# Learning the cellular origins of cancer using single-cell chromatin landscapes

**DOI:** 10.1101/2025.04.06.646131

**Authors:** Mohamad D. Bairakdar, Bruno Giotti, Wooseung Lee, Paula Stancl, Elvin Wagenblast, Dolores Hambardzumyan, Paz Polak, Rosa Karlic, Alexander M. Tsankov

**Affiliations:** Department of Genetics and Genomic Sciences, Icahn School of Medicine at Mount Sinai (ISMMS), New York, NY, USA; The Precision Immunology Institute, ISMMS, New York, NY, USA; The Tisch Cancer Institute, ISMMS, New York, NY, USA; Department of Biology, University of Zagreb, Zagreb, Croatia; Haystack Oncology, Quest Diagnostics, Baltimore, MD, USA

## Abstract

Deciphering the cell of origin (COO) of different cancers is critical for understanding tumor development and improving diagnostic and therapeutic strategies in oncology. Previous studies demonstrated that the COO chromatin accessibility landscape shapes the genomic distribution of cancer somatic mutations. We leveraged machine learning, 559 single-cell chromatin accessibility cellular profiles, and 2,734 whole genome sequencing patient samples to predict the COO of 36 cancer subtypes with high robustness and accuracy, confirming both the known anatomical and cellular origins of numerous human cancers, often at cell subset resolution. Importantly, our data-driven approach predicts that basal cells give rise to small cell lung cancers, challenging the traditional view of neuroendocrine COO. Our study also highlights distinct cellular trajectories during cancer development of different histological subtypes and uncovers an intermediate metaplastic state during tumorigenesis for multiple gastrointestinal cancers, which have important implications for cancer prevention, early detection, and treatment stratification.

## INTRODUCTION

Cancer development is a complex, multi-step process driven by genetic and epigenetic alterations that accumulate over time. A fundamental question in oncology is understanding the cell of origin (COO), or the cellular progenitor that leads to malignant transformation^1^. Identifying the COO is critical not only for understanding tumorigenesis but also for improving cancer prevention, early detection, risk stratification, and targeted treatments^1,2^. For example, precursor lesions for esophageal adenocarcinoma arise from metaplastic changes in esophageal epithelial cells due to chronic acid reflux. Understanding the COO in this context has led to targeted interventions, such as endoscopic surveillance and radiofrequency ablation, to prevent malignant progression^3,4^. Moreover, knowledge of the COO directly impacts early cancer detection; for instance, DNA methylation-based classifiers that trace circulating tumor DNA back to its tissue of origin can allow for non-invasive cancer screening^5^. Finally, studies in prostate cancer have demonstrated that tumors originating from basal versus luminal epithelial cells exhibit distinct molecular profiles and clinical outcomes and respond differently to androgen deprivation therapy^1,6,7^.

Vast progress has been made in understanding the COO of different cancers using genetically engineered mouse models^1^. However, it is also critical to directly study neoplastic processes using human samples that bypass limitations due to interspecies differences^8^. Recent advances in transcriptomic, genomic, and epigenomic profiling have emerged as powerful tools for tracing human cancer origins. Machine learning (ML) approaches have utilized extensive collections of normal and tumor bulk sequencing data to classify various cancer types according to their tissue of origin, with most methods relying on transcriptomic data^9–13^. While these approaches often achieve high prediction accuracy, the selected gene features have shown inconsistencies across different studies. More recently, these and other methods have utilized single-cell transcriptomics data to infer the COO of selected cancer types of interest^14–17^, but have not been scaled to predict COOs in a pan-cancer fashion. Additionally, approaches to detect the COO by modeling the relationship between normal and cancer transcriptomic data have limitations, as gene expression can be altered by the tumor microenvironment^18^, dedifferentiation^19^, and oncogenic reprogramming^19,20^, which can obscure the true cellular beginnings of a cancer.

Genetic data offers a more reliable means of tracing a human cancer’s COO, as the mutational landscape of cells is predominantly composed of somatic mutations that accumulate over the lifespan of an individual before malignant transformation occurs. In addition, we and others have demonstrated that the underlying epigenome of the normal COO shapes the genomic distribution of somatic mutations^21–23^, which tend to accumulate in closed chromatin regions that are less accessible by DNA repair mechanisms^24,25^. Our team first exploited the inherent relationship between epigenomic features (e.g., histone modifications, chromatin accessibility, DNA replication timing) of normal cells and the mutational landscape of eight cancers, detected using whole genome sequencing (WGS), to predict the corresponding COO using a Random Forest (RF) model^22,26^. Polak and colleagues^22^ also demonstrated that chromatin features from normal tissues are better predictors of the somatic mutational landscape than gene expression data. More recently, Yang et al.^27^ used extreme gradient boosting^28^ (XGBoost) for COO prediction, which improved prediction speed and accuracy compared to RF, especially for tumor types with low mutation density. However, the aforementioned studies^22,26,27^ were based on bulk tissue epigenomic data that lacks the resolution to identify the specific cellular populations that give rise to different cancers.

Recent advances in high-throughput, single-cell Assay for Transposase-Accessible Chromatin (scATAC-seq) have been used to profile millions of human fetal and adult cells and map chromatin accessibility across hundreds of cell types^29–35^. We reasoned that combining this data with the plethora of publicly available WGS^36–38^ data can provide unprecedented resolution and scale in predicting the COO of different cancers. We assembled one of the most extensive scATAC-seq datasets to date and leveraged our ML framework dubbed SCOOP–**S**ingle-cell **C**ell **O**f **O**rigin **P**redictor**–**to predict the COO of 36 cancer types (Supplementary Table 1), which to our knowledge represents the first genetic/epigenomic-based pan-cancer study of COOs at single-cell resolution.

Unlike bulk transcriptomic and epigenomic approaches, single-cell chromatin accessibility profiling allowed us to deconvolve complex tissues and identify cancer precursor cells at cell subset granularity. Our model demonstrated high accuracy and robustness, confirming known COOs while also uncovering novel insights into tumorigenesis. Unexpectedly, SCOOP challenged the long-held theory that small cell lung cancer (SCLC) arises from neuroendocrine cells, showing instead a basal COO, in agreement with a new study employing cellular lineage-tracing in SCLC genetically engineered mouse models^39^. SCOOP further uncovered metaplastic-like stomach goblet cells as the COO for five different gastrointestinal cancers, indicating convergent cellular trajectories toward tumorigenesis, which has important implications for cancer prevention and early detection screenings. In summary, our approach offers a robust, cost-effective framework enabling cancer biologists to ascertain the COO for a given cancer mutational profile via scATAC-seq of normal tissues of origin.

## RESULTS

### SCOOP provides improved, cellular resolution of COO predictions

To predict the COO for a cancer of interest, SCOOP uses as inputs one megabase pair binned single-nucleotide variant (SNV) count profiles aggregated across WGS patient samples, and similarly binned scATAC-seq aggregate profiles from a compendium of 559 normal cell subsets spanning 32 adult and 15 fetal tissue types (Fig. 1A; Supplementary Table 2). SCOOP leverages the binned scATAC-seq profiles and a ML model (XGBoost) to predict the mutation density of a given cancer (e.g., lung adenocarcinoma). It then iteratively reduces the set of scATAC-seq cell features through backward feature selection to identify the most informative cell subset (e.g., alveolar type II (AT2) cells), which represents the predicted COO (Methods). The model is trained 100 times using different train/test splits and random seeds (100 SCOOP runs; Methods) to obtain a robust COO prediction.

**Fig. 1:**
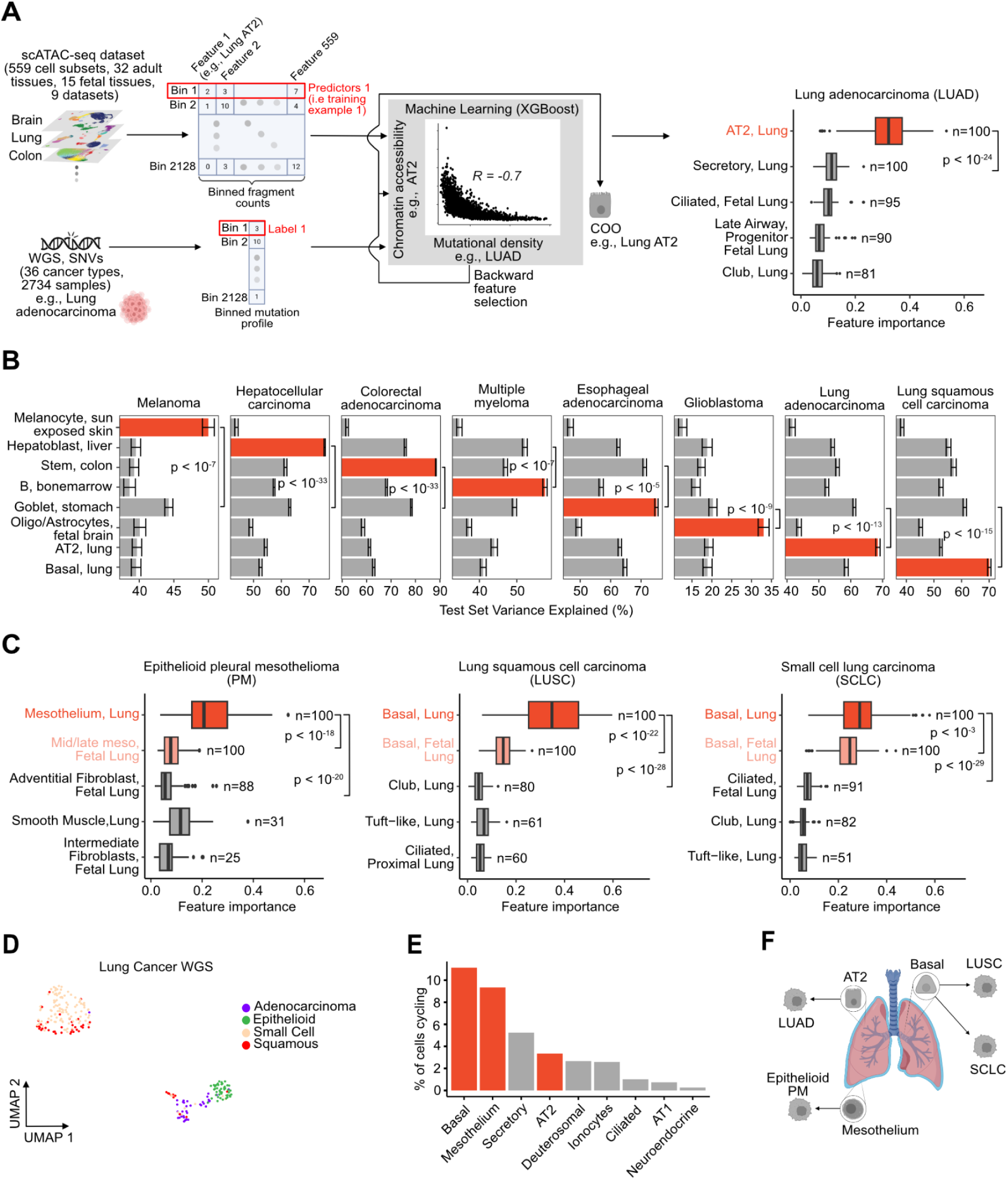
SCOOP provides improved, cellular resolution of COO predictions. **A) Left:** Illustration of how SCOOP uses scATAC-seq data to predict the COO (e.g., AT2 cells) associated with a given cancer’s mutation profile (e.g., LUAD). SCOOP takes as input a binned WGS profile of cancer SNVs and similarly binned scATAC-seq profiles from various normal cell subsets. The SNV and scATAC-seq profiles (features) are passed into a machine learning model, XGBoost, which predicts the COO through a process of backward feature selection (Methods). **Right:** Box plots of the feature importance distribution (100 SCOOP runs) of the top 5 COO predictions amongst LUAD cell subsets (Methods) (predicted COO in red). Also displayed is the number of times a cell subset appeared in the top 5 features across 100 runs (*n*). Mann-Whitney test *p*-values are displayed. **B)** Test set variance explained (*%*) by the predicted COOs (highlighted in red) for 8 cancer types studied in^22^. Error bars show the standard error of the mean (SEM) across 100 model runs. Mann-Whitney test *p*-values were computed. **C)** Box plots of the feature importance distribution (100 SCOOP runs) of the top 5 COO predictions amongst lung-related cell subsets (Methods) for PM, LUSC, and SCLC (predicted COO in red, similar cell subsets in pink). Also displayed is the number of times a cell subset appeared in the top 5 features across 100 runs (*n*). Mann-Whitney test *p*-values are displayed, where Bonferroni correction for multiple hypothesis testing was used. **D)** UMAP dimensionality reduction of individual lung cancer WGS binned mutation profiles (dots) colored by cancer type. **E)** Percentage of cycling cells across lung epithelial cell types estimated using scRNA-seq data (Methods), where COOs in E are shown in red. **F)** SCOOP’s predicted COO for different lung cancers: AT2 and mesothelial cells for LUAD and epithelioid PM respectively, and basal cells for both LUSC and SCLC. Cell type abbreviations are defined in Supplementary Table 3.

Using SCOOP, we were able to recapitulate previous tissue-level COO predictions^22^ for eight well-established cancer types but often at higher cellular granularity, given our use of single cell data rather than bulk epigenetic profiles (Fig. 1B; Supplementary Fig. 1).^22^ For instance, SCOOP predicted bone marrow B cells to give rise to multiple myeloma (MM) – which is supported by the literature^40^ – in contrast to a prior, more general hematopoietic COO prediction^22,26^. Also, melanoma was predicted to originate from melanocytes and glioblastoma (GBM) from fetal-like brain cell subsets, in agreement with previous bulk predictions^22,26^ and expected COOs^41–43^. Interestingly, our model suggested hepatoblasts as the COO for hepatocellular carcinoma (HCC), a type of hepatic progenitor cells (HPC) capable of differentiating into mature hepatocytes, with hepatocytes ranking second (Supplementary Fig. 1). Two competing theories implicate mature hepatocytes and HPCs as the source of HCC^44^, and our analysis adds weight to the conjecture that hepatoblast-like HPCs might be the primary COO of HCC. Finally, for colorectal adenocarcinoma (CRC), esophageal adenocarcinoma, lung adenocarcinoma (LUAD), and lung squamous cell carcinoma (LUSC), SCOOP pinpointed the specific cell type implicated in tumorigenesis^45–47^ – colon stem cells, mucosal stomach goblet cells, lung AT2 cells, and lung basal cells, respectively – again providing higher accuracy and cellular granularity than previous tissue-level predictions – large intestine mucosa, stomach mucosa, breast epithelial, and breast epithelial, respectively^22,26^.

### SCOOP uncovers a basal COO for small cell lung cancer

We next conducted an in-depth analysis of lung cancers, including two new types which have not been considered in previous works: pleural mesothelioma (PM) and small cell lung cancer (SCLC). For this analysis, we also added a fetal lung scATAC-seq dataset^48^ and restricted the feature space to only include lung cell subsets (Methods). To visualize SCOOP’s reproducibility, we displayed the number of appearances (*n*) and the feature importance of the 5 most informative cell subsets after backward feature selection following 100 model runs (Fig. 1A,C; Methods). SCOOP accurately and robustly predicted AT2 and basal cells as the COO of LUAD and LUSC^47^, respectively, showing significantly higher feature importance than the second most predictive feature (Fig. 1A,C; Mann-Whitney test, *p* < 10^−22^). Additionally, SCOOP’s prediction of mesothelial cells as the COO of epithelioid PM is in line with current models of mesothelial oncogenesis^49^.

To our surprise, SCOOP’s prediction for SCLC COO (Fig. 1C; Supplementary Fig. 2A when training with cell features across tissues) – adult basal cells – challenged the prevalent theory^47^ that SCLC arises primarily from pulmonary neuroendocrine cells (PNECs), also present in our feature set. Our findings are further bolstered by a previous study showing that inactivation of tumor suppressors *Rb1*, *Pten*, and *Tp53* in *Rbl1*-null murine basal cells can give rise to SCLC^50^. Additionally, we observed that LUSC and SCLC patients’ mutational density profiles clustered together and separately from LUAD and PM patients’ subclusters (Fig. 1D; Methods). This suggests an intrinsic similarity in the somatic mutational landscapes of LUSC and SCLC and further supports a shared basal COO. Of note, the LUSC and SCLC WGS data sets used by SCOOP had distinctly different genetic drivers, including an expected enrichment for *RB1* and mutations in SCLC and *NFE2L2*, *CDKN2A*, and *PIK3CA* mutations in LUSC cases, respectively (Supplementary Fig. 2B). Since cell proliferation is a hallmark of cancer, we next quantified the fraction of cycling cells across lung epithelial cell types at homeostasis using scRNA-seq data from healthy lung donors^33^ (Methods). We observed that AT2, mesothelial, and basal cells all have substantially higher proliferation rates compared to PNECs (Fig. 1E), further supporting SCOOP’s COO predictions. Taken together, our data-driven approach agrees with the accepted COO for mesothelioma, LUAD, and LUSC and also provides the first human-relevant evidence for a basal cellular origin in SCLC (Fig. 1F).

### SCOOP achieves cell subset granularity in COO predictions

We next examined SCOOP’s ability to discern the COO within intestinal and hematopoietic regenerative lineages, focusing on microsatellite stable (MSS) colorectal cancer (CRC), chronic lymphocytic leukemia (CLL), and acute myeloid leukemia (AML). CRC tumors are typically classified into 2 major subtypes based on their microsatellite status: MSS and microsatellite instable (MSI)^51^. Correlating aggregate mutational density of 51 MSS CRC^36^ with normal colon scATAC-seq data meta-cells^32^ (Fig. 2A; Methods), we observed the highest association with intestinal epithelial stem cells, the apex of the gut regenerative hierarchy. In agreement with this analysis and prior knowledge^45^, SCOOP also identifies colon stem cells as the most informative feature and COO of MSS CRC when trained using normal cell subsets across tissues (Supplementary Fig. 3) or just colon scATAC-seq data^29,30,32^ (Fig. 2B).

**Fig. 2:**
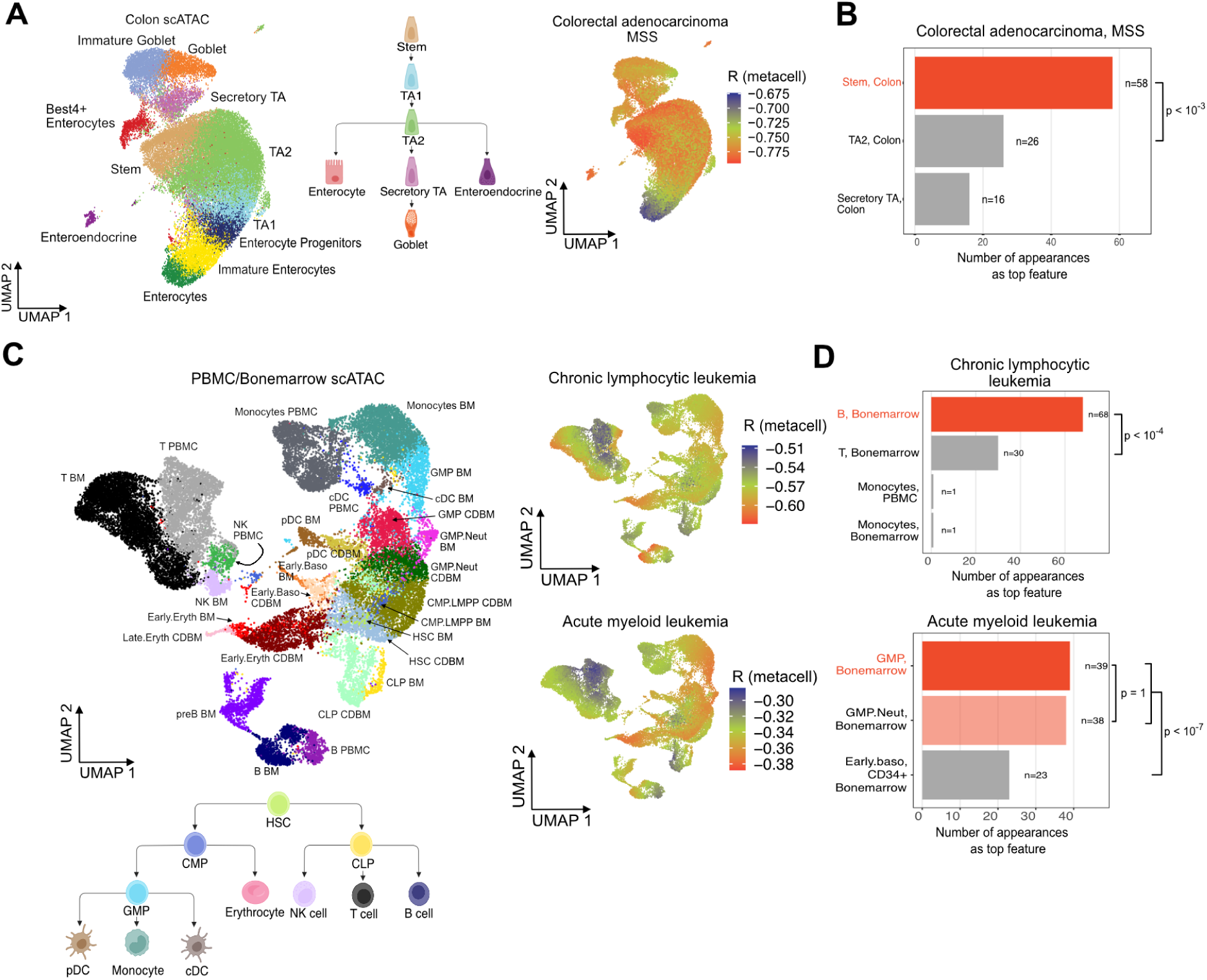
SCOOP can pinpoint cell subsets that likely give rise to different tumors. **A) Left:** UMAP of normal colon scATAC-seq data^32^ colored by cell annotations. **Middle:** Intestinal epithelial cell regenerative hierarchy. **Right:** Visualization of the Pearson correlation coefficient (R) between aggregated MSS CRC SNV profile and scATAC-seq meta-cells (Methods). **B)** Barplots of the number of times a cell subset appeared as the top feature across 100 SCOOP runs trained on normal colon scATAC-seq data^29,30,32^ and aggregated MSS CRC SNV profiles, with the predicted COO highlighted in red. Exact binomial test p-values are shown (Methods) **C) Top Left:** UMAP of all blood and bone marrow scATAC-seq cell subsets from^34^. **Bottom Left:** Hematopoietic regenerative hierarchy. **Right:** Visualization of the Pearson correlation coefficient (R) between aggregated CLL and AML SNV profiles and scATAC-seq meta-cells. **D)** Barplots of the number of times cell subsets appeared as the top feature across 100 SCOOP runs trained on scATAC-seq data from^34^ and CLL and AML aggregated SNV profiles, with the predicted COO highlighted in red and similar cell subset in pink. Exact binomial test p-values are shown (Methods), with Bonferroni correction for multiple hypothesis testing in the case of AML. Cell type abbreviations are listed in Supplementary Table 2.

We next leveraged scATAC-seq data from bone marrow and peripheral blood mononuclear cells (PBMCs)^34^ to investigate the COO of CLL and AML (Fig. 2C-D)^52^. Meta-cell correlation analysis showed that CLL was most anti-correlated with bone marrow B cells, but not PBMC B cells (Fig. 2C; top right). In agreement, SCOOP predicts bone marrow B cells as the COO (Fig. 2D; Supplementary Fig. 3 when training across tissues), supporting the prevailing hypotheses that B cells give rise to CLL^53^. In contrast, AML exhibited less specificity in terms of correlation patterns, possibly due to the high interpatient heterogeneity observed in AML^54^, being highly anti-correlated with multiple cell types, most notably bone marrow granulocyte macrophage progenitors (GMP) and early erythrocytes (Early.Eryth) (Fig. 2C; bottom right). Prior work has shown leukemia stem cell (LSC) transcriptional activity in cells resembles lymphoid-primed multipotent progenitors (LMPPs) and GMPs rather than hematopoietic stem cells (HSCs), suggesting that LSC transformation largely occurs at the progenitor stage, either directly from progenitors with abnormal self-renewal capabilities or from HSCs upon further differentiation^55^. Additionally, studies using murine leukemia models and various genetic modifications indicate that both HSCs and committed myeloid progenitor cells can evolve into LSCs, which phenotypically and molecularly resemble committed myeloid progenitor cells^56,57^. Curiously, SCOOP supported these results: across 100 runs of the model, highly similar cell subsets bone marrow GMP and GMP/Neutrophils (GMP.Neut) – types of myeloid progenitors – appeared a combined 77 times as the top predicted feature (Fig. 2D). This bolsters the hypothesis that myeloid lineage differentiation is a prerequisite for AML development^58^. Finally, we note that comparing our meta-cell correlation analysis to our machine learning-based predictions highlights SCOOP’s capacity to exploit non-linear relationships that may not be easily captured by linear correlation analyses.

### Distinct COOs for cancers of different histologies

Besides displaying cell subset granularity, SCOOP also identified histological cancer subtypes with different COOs. We first examined three subtypes of kidney renal cell carcinoma (RCC): clear cell RCC (ccRCC), papillary RCC (pRCC), and chromophobe RCC (chRCC). Both ccRCC and pRCC are thought to originate from the proximal tubule in the kidney, while it is suspected that chRCC originates from the distal tubule^59^. In agreement, individual patient somatic mutation profiles demonstrate higher similarity between ccRCC and pRCC in comparison to chRCC (Fig. 3A). SCOOP also correctly matched ccRCC and pRCC COO to proximal tubule progenitor-like (PTPL) cells, again demonstrating cell subset resolution, while identifying chRCC COO as collecting duct, intercalated cell type A (ICA) from the distal tubule (Fig. 3B; Supplementary Fig. 4 when training with cell features across tissues).

**Fig. 3:**
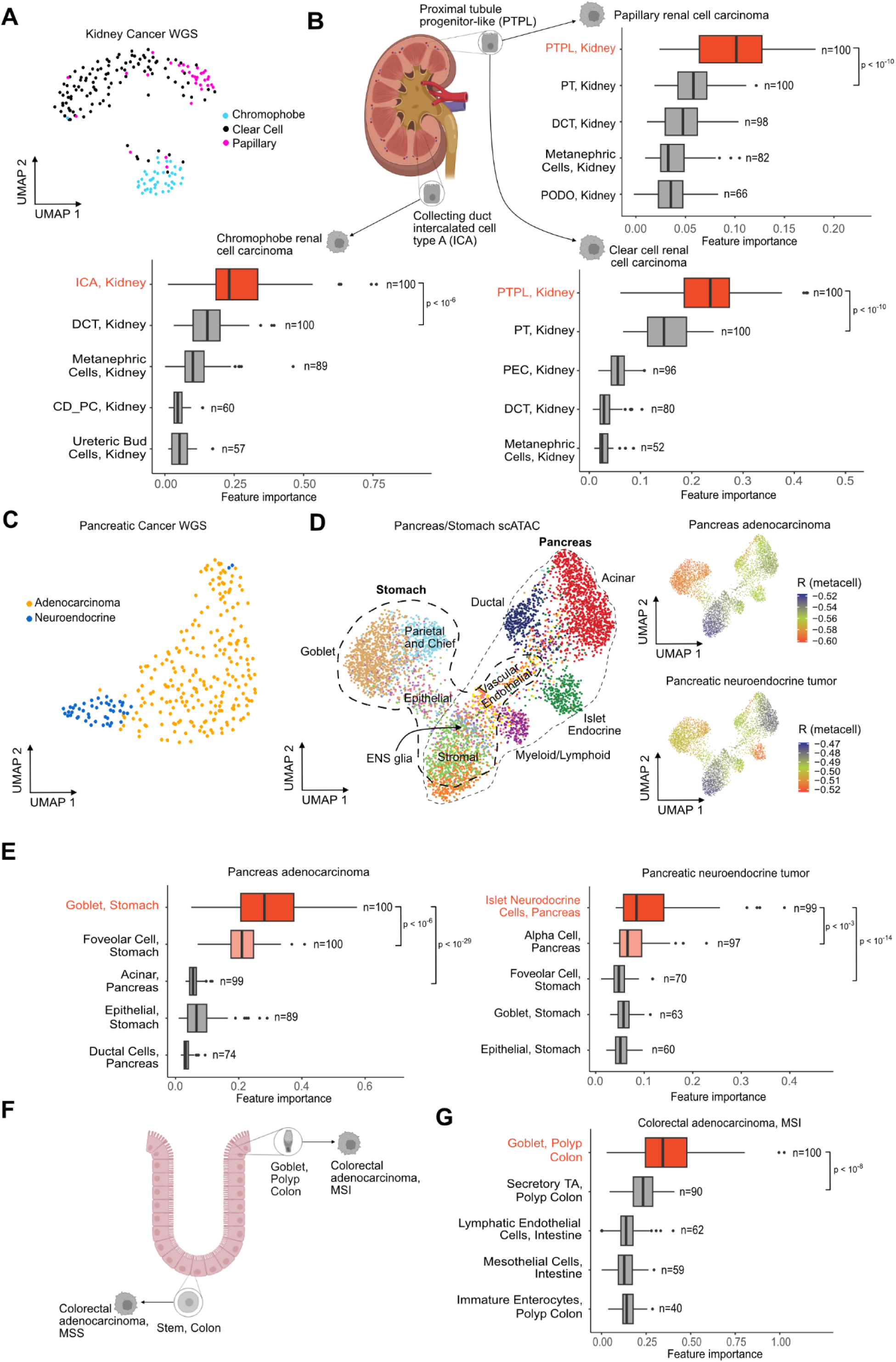
Histological cancer subtypes are associated with different COOs. **A)** UMAP of individual kidney cancer patient binned WGS mutational profiles from three different kidney cancer subtypes: ccRCC, pRCC, and chRCC **B)** Box plots of the feature importance distribution (100 SCOOP runs) of the top 5 COO predictions amongst kidney-related cell subsets (Methods) for ccRCC, pRCC, and chRCC (predicted COO in red). Also displayed is the number of times the feature appeared in the top 5 features across the 100 runs (n). Mann-Whitney test *p*-values were computed. **C)** UMAP dimensionality reduction of individual pancreatic cancer WGS samples binned mutation profiles (dots) colored by cancer type. **D) Left:** UMAP of stomach and pancreas scATAC-seq data from^30^. Thinner dashed line encompasses pancreatic cells, whereas thicker dashed line demarcates stomach cells. **Right:** UMAPs displaying the Pearson correlation coefficient (R) between aggregated PAAD and PNET mutational profiles and scATAC-seq meta-cells (Methods). **E)** Box plots of the feature importance distribution (100 SCOOP runs) of the top 5 COO predictions amongst pancreas- and stomach-related cell subsets (Methods) for PAAD and PNET (predicted COOs highlighted in red, similar cell subsets in pink). Mann-Whitney test *p*-values were computed, with Bonferroni correction for multiple hypothesis testing. **F)** Accepted model of CRC COOs agrees with SCOOP’s predictions: colon goblet cells for MSI, and stem cells for MSS. **G)** Box plot of the feature importance distribution (100 SCOOP runs) of the top 5 COO predictions amongst colon-related cell subsets (Methods) for CRC, MSI (predicted COO in red). Mann-Whitney test *p*-values were computed. Cell type abbreviations are defined in Supplementary Table 2.

Exploring the cellular origins of pancreatic adenocarcinoma (PAAD) and pancreatic neuroendocrine tumor (PNET), we again observed intrinsic differences in tumors’ mutational profiles that cluster by histological subtypes (Fig. 3C). It is posited that the genesis of PAAD is rooted in a transformation from acinar to ductal cells, known as acinar-to-ductal metaplasia (ADM)^60^, while PNETs are thought to arise from islet cells, which are part of the endocrine system of the pancreas^61^. In agreement, we observed that PAAD and PNET aggregated mutational profiles were most anti-correlated with stomach goblet cells and islet endocrine cells (Fig. 3D), respectively, and SCOOP further reinforced these results when trained using all pancreas and stomach cell features (Fig. 3E). Goblet cells secrete mucus to protect the stomach lining, which matches previous bulk-level prediction of stomach mucosa and suggests metaplasia transformation in PAAD^26^. Our PNET COO prediction of pancreatic islet endocrine cells was also in agreement with PNET bulk-tissue prediction^26^, but came in second when training SCOOP across different tissues to the highly similar colon endocrine cells (Supplementary Fig. 4), highlighting the benefit of restricting SCOOP’s feature space to anatomically relevant cell subsets based on prior knowledge.

Motivated by previous studies arguing that the MSS CRC subtype arises from stem cells in the colon crypt, while the MSI subtype arises from gastric metaplasia^62^, we wanted to investigate this hypothesis (Fig. 3F) in more detail. To capture the metaplasia cell state relevant for MSI tumor COO, we added precancerous polyp scATAC-seq data^32^ to our feature space. SCOOP supported the hypothesis that MSI tumors likely arise from a metaplasia cell state distinct from MSS CRC tumorigenesis trajectory and further pointed specifically to polyp colon goblet cells as the COO (Fig. 3G; Supplementary Fig. 4).

### Gliomas likely arise from fetal-like multipotent progenitor cells

Inspired by SCOOP’s success in accurately linking distinct histological subtypes with their COO, we next examined the genesis of different brain cancers. In light of prior research indicating a significant role of fetal-like neural stem cells (NSCs) and oligodendrocyte progenitor cells (OPCs) in brain tumorigenesis^42,63,64^, we leveraged two additional scATAC-seq datasets that characterized fetal and adult brain cell subsets’ chromatin accessibility extensively^31,65^. Interestingly, SCOOP predicted that medulloblastoma (MB), GBM, pilocytic astrocytoma (PA), and oligodendroglioma (OG) all originate from fetal-like cell subsets (Fig. 4A; Supplementary Fig. 5 when training with cell features across tissues), in agreement with previous bulk epigenomic data modeling^22,26^. For MB, SCOOP predicted granule neurons from fetal cerebellum as the COO, closely matching the anatomical and cellular origin (cerebellar granule neuron precursor cells) for one of the four major subtypes of MB, the Sonic Hedgehog (SHH) subtype^63^. The other major subtypes – Group 3, Group 4, and WNT – are thought to arise from unipolar brush cells (UBCs), neural stem, or progenitor cells, all of which were not present in our scATAC-seq dataset^66^.

**Fig. 4:**
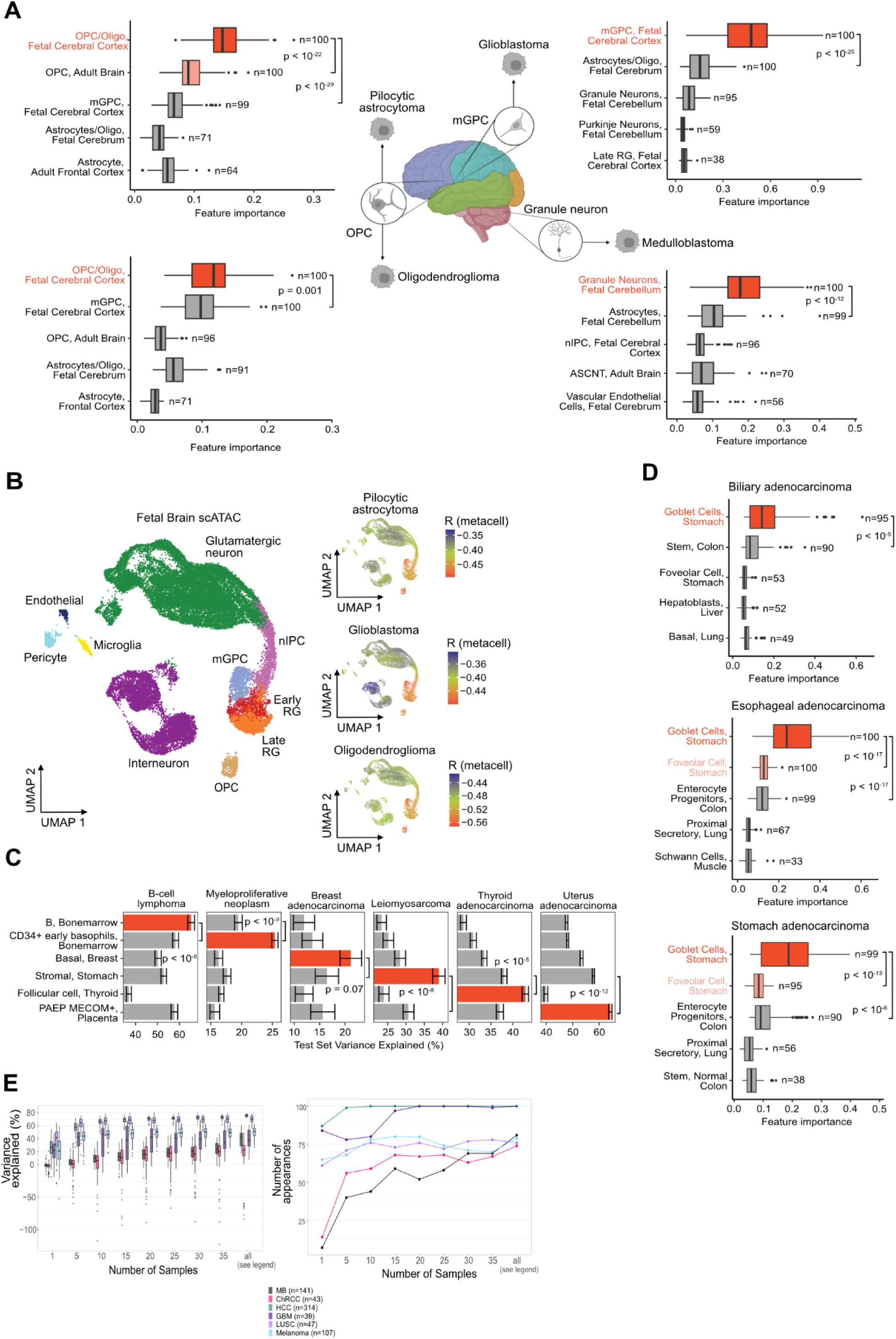
Pan-cancer COO predictions across 36 cancer types. **A)** Box plots of the feature importance distribution (100 SCOOP runs) of the top 5 COO predictions amongst brain-related cell subsets (Methods) for GBM, OG, PA, and MB (predicted COO highlighted in red, similar cell subsets in pink). Also displayed is the number of times the feature appeared in the top 5 features across the 100 runs (n). Mann-Whitney test *p*-values were computed, with Bonferroni correction for multiple hypothesis testing in the case of OG. **B) Left:** UMAP of fetal brain scATAC-seq data cell subsets from^31^. **Right:** UMAPs displaying the Pearson correlation coefficient (R) between aggregated PA, GBM, and OG mutational profiles and scATAC-seq meta-cells. **C)** Test set variance explained (%) by the predicted COOs across 6 additional cancer types. The predicted COO for each cancer type is highlighted in red (also see Supplementary Fig. 6). Error bars show the standard error of the mean (SEM), after 100 SCOOP runs. Mann-Whitney test *p*-values were computed. **D)** Box plots of the feature importance distribution (100 SCOOP runs) of the top 5 COO predictions for biliary, esophageal, and stomach cancer (predicted COO highlighted in red, similar cell subsets or alternative COO predictions in pink). Mann-Whitney test *p*-values were computed, with Bonferroni correction for multiple hypothesis testing in the case of stomach and esophageal adenocarcinoma **E)** Model performance as a function of WGS sample size for tumors with high (LUSC, melanoma), moderate (HCC, GBM), and low (ChRCC, MB) TMB tumors. **Left:** Distribution of variance explained across 100 model runs for different number of WGS samples, randomly subsampled. **Right:** Number of appearances of the expected COO as the top feature across 100 model runs for different WGS sample sizes. Cell type abbreviations are listed in Supplementary Table 2.

For both pilocytic astrocytoma (PA) and oligodendroglioma (OG) SCOOP predicted the COO to be OPCs from the fetal cerebral cortex. Despite its namesake, recent studies have suggested that PA originates from the oligodendrocyte lineage^67^. Also, PA cancer cells were compared with various brain cell types using single cell transcriptomic atlases, and it was found that they exhibited a gene expression signature most similar to OPCs^68^. Finally, for GBM, while SCOOP originally predicted Oligo/Astrocytes from the fetal brain (Fig. 1B) as the COO, after adding the comprehensive fetal brain dataset, the COO was more specifically predicted as multipotent glial progenitor cells (mGPCs) from the fetal cerebral cortex. This is further supported by our recent fetal brain cell atlas, showing high transcriptional similarity of GBM malignant cells to multipotent progenitor fetal cell populations^69^. We observed highly similar cell subset specificity when conducting a meta-cell correlation analysis on scATAC-seq data from^31^ (Fig. 4B). One limitation of our glioma findings, is that our model does not contain adult NSC epigenomes, which have been implicated in the genesis of adult GBM and OG^43^. However, our datasets contain fetal multipotent progenitor cells (mGPC, nIPC, radial glial cells) that can give rise to different glial and neuronal populations akin to NSCs^64^.

### Pan-cancer predictions across 36 cancer types identifies metaplasia-mediated neoplasms

We next applied SCOOP to WGS data from B-cell lymphoma, myeloproliferative neoplasm (MPN), breast adenocarcinoma, leiomyosarcoma, thyroid, and endometrial cancer, where each prediction matched one of the putative COOs^70–73^ (Fig. 4C; Supplementary Fig. 6A). In the case of leiomyosarcoma, SCOOP predicted stromal cells as the COO, which show high chromatin accessibility at established smooth muscle cell marker loci (Supplementary Fig. 6B). For endometrial cancer, the predicted COO – PAEP/MECOM positive cells from the placenta – corresponds to endometrial epithelial cells^30^. MPN most likely arises from HSCs^74^ but there exists evidence that it may also arise from committed hematopoietic progenitors similar to HSCs^75^. SCOOP’s top COO predictions for MPN are bone marrow CD34+ early basophils, GMP, and CMP.LMPP (Supplementary Fig. 6A, when using tissue-specific and all features), all three representing multipotent hematopoietic progenitors with close proximity to HSCs in their epigenome and regenerative hierarchy. Worth noting, MPN has both the lowest total number of mutations (*n*=18,512) and average mutations per sample (*µ*=805) among the cancer types we analyzed (Supplementary Table 1), which may explain the variance in its prediction. Finally, while SCOOP predicted basal epithelial cells from mammary tissue as the COO for breast adenocarcinoma – as opposed to luminal cells (present in our dataset) which are considered a more predominant COO – basal cells still constitute a possible COO^26,76^.

Similar to PAAD, three other gastrointestinal cancers – biliary, esophageal, and stomach adenocarcinoma – were predicted to arise from stomach goblet cells (Fig. 4D), which matches previous bulk-level predictions of stomach mucosa and also suggests metaplasia transformation in these cancers^26^. In agreement, esophageal adenocarcinoma formation is thought to undergo a transitional metaplasia phase, called Barrett’s esophagus^46^. While previous modeling has implicated liver (33.3%), mucosa (∼25%), and breast epithelial (∼20%) bulk epigenomes in the genesis of different biliary tract cancers^77^, our cell type specific results argue that at least a subset of biliary cancers undergo metaplasia.

We also report COO predictions for other cancer types with available WGS data (Supplementary Fig. 7), which we categorized as either 1) matching a proxy cell type, 2) missing expected COO scATAC-seq data, or 3) demonstrating low variance explained (<10%). Category 1 included cervical and head and neck squamous cell carcinoma COO predictions of esophageal epithelial cells, which was the closest comparative cell profile in our dataset and in^26^ to the expected COO for these tumors. Osteosarcoma also fell into Category 1, matching stromal cells from the heart, as it is expected to arise from mesenchymal cells^78^. Category 2 included prostate cancer, and ovarian cancer, and bladder transitional cell carcinoma (Supplementary Fig. 7). Category 3 consisted of breast lobular carcinoma, for which the median variance explained of the top feature was lower than 10% (Supplementary Fig. 7).

We also quantified the relationship between WGS sample size and SCOOP’s prediction accuracy for cancers with low, medium, and high average tumor mutational burden (TMB; Supplementary Table 1; Methods). Interestingly, for both medium and high TMB cancers examined, as few as 5 WGS samples were enough for SCOOP to identify the correct COO at least 68% of runs (≥42.4% variance explained), while low TMB tumors (e.g., ChRCC, MB) required 30 samples to achieve similar levels of performance (correct COO ≥69% of runs; Fig. 4E).

## DISCUSSION

Our work combined machine learning, WGS, and scATAC-seq data to predict the COO of 36 human cancer subtypes. Our approach employing single-cell epigenomic data marks a significant advancement, offering unparalleled resolution in the cellular subsets, developmental, and regenerative hierarchies underlying the genesis of cancer. For most cancers examined in our study, the predicted COOs and anatomical location were highly reproducible and aligned closely with those posited by prior research, serving as a validation of both the existing scientific consensus and the accuracy and reliability of our methodology.

Beyond validation, the true potential of our approach lies in its capacity for data-driven hypothesis generation, particularly for cancers with ambiguous or unknown cellular and anatomical origins. Our study’s surprising finding of a basal COO for SCLC, which has historically been thought to arise from neuroendocrine cells based on histological and transcriptomic similarities^47^, exemplified the potential of our ML approach to uncover novel avenues for research. Substantiating our computational approach, a recent study employing lineage tracing in genetically engineered mouse models of SCLC supported the prediction that basal cells may constitute the predominant COO in SCLC as they can give rise to all the SCLC subtypes upon transformation and in proportions matching human epidemiological data^39^. These findings have important clinical implications for prevention, early detection, and treatment of SCLC. For example, early screening programs and smoking cessation strategies that target individuals with molecular changes in bronchial basal cells^79^ would be highly relevant to not just lung squamous cell carcinoma but also SCLC tumors.

While SCOOP uncovered basal cells as the predominant COO of SCLC, future research will be required to investigate if this is the case for all SCLC subtypes and, more generally, for neuroendocrine tumors (NETs) across tissues. For example, olfactory neuroblastomas^80^ display subtype heterogeneity resembling that of SCLC and were shown to arise from globose basal cells using genetically engineered mouse models^80^. In contrast, pulmonary carcinoids are less aggressive NETs that may arise from a different COO, given that they are not associated with smoking, occur in younger individuals, and are considered as a separate class of tumors from SCLC^81,82^. Moreover, we recently identified a highly proliferative, tuft-ionocyte progenitor (TIP) cell^33^ in the human lung, which appeared among the top five most predictive features for SCLC; it is tempting to speculate that TIP cells may also play a role in a subset of SCLC transformations, especially for the rare tuft-like, POU2F3+ SCLC tumors.

Our study also identified stomach goblet cells as the most predictive epigenomic feature for the somatic mutational landscape of several cancer types–PDAC, MSI CRC, biliary, esophageal, and stomach adenocarcinoma. These results are compatible with an intermediate metaplastic state contributing to tumorigenesis in multiple cancers across diverse tissues and organs, and is further corroborated by prior bulk-level model predictions^26^. Metaplasia, a process in which one differentiated cell type is replaced by another, is a well-known precursor to cancers such as esophageal adenocarcinoma and gastric cancer^46,83^. Our findings suggest that metaplastic transitions for different cancers may be more widespread than previously recognized, which can inform the development of new metaplasia biomarkers, akin to stomach goblet cells, for improving early detection across multiple cancers. Furthermore, pan-cancer identification of tumors that arise from metaplasia can result in sharing and repurposing of successful prevention^84^, risk assessment, and treatment strategies across metaplasia-related cancers.

Despite the unprecedented resolution, accuracy, and scale of SCOOP’s COO predictions, our ML approach has several limitations that will require future research. A fundamental impediment arises from previous observations^22,26^ that robust model prediction requires aggregation of WGS profiles from different tumor samples that may have different COOs. This presents challenges for obtaining personalized predictions, which can be remedied in the future by acquiring additional WGS data that can enable grouping of patient tumors with similar COOs. Another limitation pertains to the comprehensiveness of our single-cell atlas and data quality across tissues, potentially omitting relevant cell subsets that could serve as the COO for specific cancers. Finally, SCOOP identifies the pre-malignant cellular ancestor whose chromatin accessibility best explains a cancer’s cumulative somatic mutational landscape. In some cases, this cell type can differ from the normal cell that acquires the initial oncogenic hit, as appears to be the case for PDAC, other metaplasia-related cancers in our study, AML, and as shown previously in gliomas^42,43^ and other cancers; future modeling and experimental studies tracing the transition from precancerous to malignant cell states will be necessary to uncover the exact trajectory of cellular transitions involved in tumorigenesis.

SCOOP contrasts with previous approaches that necessitated individual experiments for each sample—for instance, conducting 100 separate experiments to generate 100 ATAC-seq profiles from bulk tissue. Instead, using publicly available scATAC-seq data, our approach successfully derived 559 distinct cell subset profiles from 42 tissue experiments. This enhanced efficiency not only facilitates the identification of a wider variety of cell types, but also paves the way for a cost-effective framework for COO identification by substantially decreasing the need for additional sequencing experiments, streamlining laboratory procedures. Future studies will also benefit from the increasing number and quality of WGS and scATAC-seq data being generated, and employ SCOOP to investigate rare cancers and more refined histological and molecular subtypes not examined here. Moreover, expanded scATAC-seq sampling across the human body can further enhance our method’s ability to identify the anatomical location for a cancer’s COO, as demonstrated for MB (cerebellum). Our easy-to-use computational platform is accessible to all cancer biologists, requiring only the availability of WGS data for a cancer of interest and, if necessary, scATAC-seq data from the corresponding normal tissue to complement our 559 normal cell subsets already assembled.

## MATERIALS AND METHODS

### Whole genome sequencing (WGS) data acquisition and processing

All cancer WGS data besides that for mesothelioma, SCLC, multiple myeloma, and MSI-high CRC were obtained from the Pan-Cancer Analysis of Whole Genomes (PCAWG) study^36^. PCAWG samples were obtained both from the publicly available, International Cancer Genome Consortium (ICGC) portion of PCAWG via the ICGC data portal^36^, and from the restricted access TCGA portion of PCAWG via Bionimbus^85^. Only samples on the tumor whitelist from PCAWG were processed for analysis. For pleural mesothelioma, the somatic mutation genome locations were acquired directly from the authors of ^38^. We restricted our analysis to epithelioid pleural mesothelioma, since the number of samples for the other two histological subtypes (sarcomatoid and biphasic) was very low (2 and 3 respectively). We also excluded samples that were classified as “Not Otherwise Specified.” For SCLC, the SNV genomic locations were obtained from the authors of George et al.^37^ through the European Genome-Phenome Archive (EGA). Multiple myeloma data was obtained and processed as described previously^22^. Finally, MSI-high data was obtained from TCGA. We obtained single-nucleotide variants (SNVs) from WGS data for each cancer type examined in our study and aggregated the variant counts across samples into 1 megabase pair bins, excluding sex chromosomes, as we described previously in^22^. In brief, all somatic single nucleotide mutation data per cohort were converted to BED format, and intersected using BEDtools with the 1MB bins. The number of mutations per bin were then aggregated by cancer type/subtype. As explained in^22^, these bins exclude regions that overlap centromeres and telomeres, and regions where the fraction of mappable base pairs is lower than 0.92.

### Somatic variant calling for MSI CRC WGS samples

For the microsatellite instability (MSI) samples, both tumor and matched normal BAM files containing mapped reads onto the human version hg38 were downloaded. We applied an ensemble consensus variant calling approach utilizing Mutect2^86^ (GATK v4.3.0.0), Strelka2^87^ (v2.9.10), and VarScan^88^ (v2.4.6), retaining only SNPs identified by at least two callers. Mutect2 analysis included the use of The Panel of Normals (PoN) and germline resources following GATK Best Practices. Subsequently, the filtered SNPs were converted to hg19 using the R package liftOver (v1.26.0).

### scATAC-seq data acquisition and processing

scATAC-seq data for all cell subsets used in this study were obtained from multiple previously published scATAC-seq datasets. 222 fetal and adult cell subsets from 30 adult and 15 fetal tissues were obtained from scATAC-seq atlases in^29,30^. More specialized scATAC-seq data from adult brain^65^, blood and bone marrow^34^, colon^32^, lung^33^, and kidney^35^, as well as fetal lung^48^ and brain^31^, were also included in the final dataset. We note that data from separate datasets were kept separate and were not merged. For all datasets except kidney^35^, fragment files were available and were migrated to hg19 if they were not already aligned to hg19 using the R package liftOver (v1.26.0). For kidney scATAC-seq data, we aligned FASTQ files to hg19 and obtained fragment files using Cell Ranger ATAC pipeline (v1.1.0). scATAC-seq data fragment counts were binned as described above for SNV counts from WGS data; in brief, scATAC-seq fragment counts across cells were aggregated into bins to obtain chromatin accessibility profiles for each cell subset. Each cell subset (feature) is followed by a dataset identifier: D1 for ^29^, D2 for ^30^, D3 for ^32^, D4 for ^34^, D5 for ^33^, D6 for ^35^, D7 for ^31^, D8 for ^65^, and D9 for ^48^.

### scATAC-seq data curation and annotation

scATAC-seq cell annotations for each individual dataset were obtained directly from the corresponding paper’s published materials, except for the lung dataset^33^ for which the annotations were not available at the time of writing, and for which we generated de novo annotations. After reviewing the publicly available annotations, we further refined them as follows: for the adult atlas^29^, brain^65^, blood and bone marrow^34^, kidney^35^, all cell types that had a number index at the end indicating cell-type sub-clusters were collapsed into a single combined cell type. For example, “Colon Epithelial Cell 1”, “Colon Epithelial Cell 2”, and “Colon Epithelial Cell 3” from Transverse Colon were annotated as “Colon Epithelial Cell”. For the blood and bone marrow dataset^34^, we additionally removed cell types annotated as unknown, and combined all T-cell subsets (i.e all CD8 and CD4 subsets) into T cells. For the lung dataset^33^, we used an analogous approach to that used in the original publication. In particular, the cell type annotation was informed by labels from scRNA-seq data and was implemented in ArchR. These labels were transferred to the scATAC-seq data using the *addGeneIntegrationMatrix* function. Marker discovery was conducted de novo using the *getMarkerFeatures* function with the GeneScoreMatrix assay. The final annotation of scATAC-seq cells was carried out by mapping the newly discovered clusters to predicted RNA cell types, or, when no corresponding cell type was found in the RNA data, by linking them to the chromatin accessibility at the locus of known gene markers that were most prevalent in each cluster. For the fetal atlas^30^, we excluded all cell types that were annotated as unknown. For the fetal brain dataset, we used the same annotations as displayed in the scATAC-seq UMAP of the original paper (see Fig. 1F of ^31^), except we removed cell types annotated as unknown. We did not use cancer tissue scATAC-seq data from ^32^; specifically, our compiled dataset included only normal and unaffected samples, both of which were considered as “normal” tissue. Except for the five blood cancers (MM, CLL, AML, BNHL, and MPN), where it was important to distinguish between CD34+ and CD34-bone marrow, we collapsed the annotation into a single bone marrow category (Supplementary Table 2 contains the uncollapsed numbers). For MSI CRC modeling that is expected to undergo metaplasia, our modeling also included polyp samples that are expected to contain this stage in tumor development. No annotation curation was done for the fetal lung dataset^48^.

After finalizing our annotations, we applied a filter of a minimum 100 cells per feature to exclude cell subsets with insufficient number of fragments to produce an accurate pseudo-bulk epigenetic profile, as quantified in^89^. If a given cell type did not meet this threshold, it was excluded from further analysis, except for adult lung neuroendocrine cells from^33^ when conducting lung-specific analyses, since they were conjectured to be the COO of SCLC. While our dataset contained only 41 adult PNECs, it also included 356 GHRL+ and 231 fetal PNECs, which is comparable to the number of mesothelial cells (n=401 and 223 for adult and fetal).

### Feature space selection for modeling

We trained our model on a variety of different feature spaces. Except when indicated otherwise, we trained our models using 6 different datasets: two scATAC-seq atlases^29,30^, blood and bone marrow^34^, colon^32^, lung^33^, and kidney^35^. The other datasets were included in the analysis when we dove deeper into a select cancer type and interrogated its COO more comprehensively.

### Cancer subtype classification

We used the metadata in cBioPortal^90^ to obtain MSI high vs MSS classification for CRC. In particular, if the metadata column “subtype” had “MSI” as a suffix, we considered the sample to be MSI high. Otherwise, it was considered to be MSS. We note that all samples for ColoRect-AdenoCA from PCAWG are MSS samples, except for one, which is MSI high. We used the metadata in ICGC^36^ to obtain pRCC and ccRCC classification for Kidney cancer.

### XGBoost regression model

We trained XGBoost (Extreme Gradient Boosting) regression models using XGBRegressor from the Python XGBoost package version 1.7.4. XGBoost is an advanced implementation of the gradient boosting algorithm, designed for speed and performance. It builds an ensemble of decision trees in a sequential manner, where each tree corrects the errors of its predecessor. XGBoost employs a regularized model formalization to control over-fitting, making it robust to noisy data. The algorithm is parallelizable across both cores in a CPU and machines in a distributed setting, resulting in significantly faster training times compared to traditional gradient boosting. This enabled us to perform robustness analysis by performing 100 runs for each model.

We frame our task as a machine learning regression problem as follows: each genomic bin corresponds to a training example (i.e data point) where the input features (i.e “predictors” or “independent variables”) are the aggregated scATAC counts from the various cell subsets for that bin, and the label (i.e “response variable” or “target variable” or “dependent variable”) is the mutation count for that bin. Put differently, each genomic bin is characterized by two sets of numbers; one set (input) is composed of multiple “features” where a particular “feature” constitutes an aggregated scATAC count for a particular cell subset, while the other set (output) is comprised of a single number: the aggregated number of mutations within that bin.

#### Robustness analysis

After partitioning the genome into bins, we grouped contiguous bins into 10 train/test folds. We used each fold as a test fold 10 times and used the remaining 9 folds for training and validation using cross-validation with a 90/10% split of training and validation respectively. For each of the 10 runs of cross validation and model testing per fold, 10 different seeds were used for seeding the XGBoost model building process, and, when training models with more than one feature, the feature importance calculation. Since we used each of the 10 folds as a test set 10 times (with a different seed each time), we in effect obtained 100 estimates of model performance on an unbiased test set.

#### Feature selection

If the initial feature space had more than 20 features, we next selected the top 20 features according to our feature importance score. Otherwise, the entire feature space was used to proceed. Using this reduced (or unmodified) feature space, we then performed iterative backward feature selection until only a single feature remained, which in most cases we would expect to correspond to the cell of origin for the cancer type under consideration (see Robustness box plots and COO prediction section for more details). To be more specific, after reducing the feature space to 20 features, we trained a new model using this reduced feature space, picked the best model according to mean performance across validation folds when performing cross-validation, ranked the 20 features based on the chosen model, and eliminated the bottom feature. This was then repeated iteratively.

We note that performing this process may aid in alleviating the potential bias that can be induced by having correlated features. As an example, suppose our dataset contains cell types A, B, and C, where B and C are highly correlated. When computing feature importance, cell types B and C may be ranked lower than they would have been ranked otherwise if the other cell type were absent. Thus, if the model ranks B above C and we remove cell type C, cell type B can now more fairly compete against cell type A for the top spot. There is the further issue that B and C may be arbitrarily ranked above one another by the model, which motivates training the model multiple times using different random seeds.

#### Feature importance calculation

Feature importance was calculated using the “permutation_importance” function from scikit-learn (v1.3.0), setting the “n_repeats” parameter to 10, after picking the best model according to cross-validation. This function implements a permutation importance mechanism:for each feature in the feature space, it randomly permutes its values and keeps all other features constant, then measuring the effect of this permutation on a model’s chosen performance metric by comparing the change in performance to the baseline performance when no permutation is performed. The larger the negative effect of this process on the chosen performance metric, the more important a feature is deemed to be. Since this permutation is a random process, it is useful to perform this process multiple times, and measure the mean effect of permutation on performance. We chose to repeat the process 100 times. We note that since we used cross-validation, feature importance was calculated on each validation fold and averaged across folds; since we used a 90/10% train/validation split, which implies 10 train/validation folds, and for each fold we computed 10 permutation scores (n_repeats=10), this in effect means that we performed 100 permutations and averaged these for any given feature.

#### Model evaluation metric

We assessed model performance by computing the R^2^ score (i.e variance explained) of our model. This is computed as 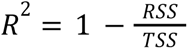, where RSS is the residual sum of squares, and TSS is the total sum of squares. We emphasize here that it is possible to obtain a negative value for *R*^2^, if the model performs worse than a simple mean model. This situation occurs when the RSS is greater than TSS, which means that the model’s predictions are on average further from the actual values than the simple mean of the data.

#### Hyperparameter optimization

Hyperparameter optimization was performed using the automated hyperparameter search framework Optuna^91^ (v3.3.0). We used the default hyperparameter optimization strategy, Tree-structured Parzen Estimator (TPE), which is a Bayesian optimization strategy. Briefly, we note that, in contrast to the naive but commonly employed strategy of randomly choosing and testing model hyper-parameters settings e.g grid search or random search – which is an inefficient and non-optimal strategy – Bayesian optimization strategies like TPE take advantage of the history of model performance under different hyperparameter settings to cleverly explore the hyperparameter search space, and exploit settings that performed well to narrow down the search space for optimal hyperparameters.

Each training run in Optuna is called a “study.” Each study consists of multiple “trials,” each corresponding to a specific model hyperparameter setting. At the end of a study, the best model hyperparameters are chosen based on the trial that performed best according to some pre-specified metric. In our case, this is the mean variance explained across validation folds. In other words, we fix a hyperparameter setting, compute its mean performance across validation folds during cross validation, report this number as the performance for the hyperparameter setting in question, and repeat this process for different hyperparameter settings. We note that the number of trials per study must be specified, and we set it to 50 (i.e the “n_trials” parameter of the “study.optimize” function is set to 50). In practice, this means that 50 hyperparameter settings are tested per training run of the model. We emphasize that during backward feature selection, this process is repeated from scratch i.e for each new feature space, 50 different hyperparameter settings are tested, and the best models chosen for different feature spaces very likely differ in their hyperparameter settings.

Table 1 below lists the XGBoost model hyperparameters and the corresponding ranges of values we searched over. The full description of each hyperparameter can be found at https://xgboost.readthedocs.io/en/release_1.7.0/python/python_api.html, under xgboost.XGBRegressor.

**Table 1:** Table showing the XGBoost hyperparameter search space that was used for Optuna, including the hyperparameter name, the type of the hyperparameter, the range of values explored, and the scale of the search (i.e, linear, log).

| Hyperparameter | Type | Range | Scale |
| --- | --- | --- | --- |
| n_estimators | integer | 100-500 | linear |
| max_depth | integer | 3-10 | linear |
| learning_rate | float | $1e^{-8}$-1.0 | log |
| subsample | float | 0.1-1.0 | linear |
| colsample_bytree | float | 0.1-1.0 | linear |
| min_child_weight | integer | 1-6 | linear |
| reg_lambda | float | $1e^{-8}$-1.0 | log |
| reg_alpha | float | $1e^{-8}$-1.0 | log |

#### Robustness box plots

For the robustness box plots (e.g Fig. 1C), we display, across 100 runs, the feature importance of different features when the model was trained on x features i.e after conducting backward feature selection and being left with x features. We further restricted the display to only show the top 5 features, where we define “top 5 features” as the features that appeared most frequently in the top 5 across 100 runs of the model. Also displayed is the number of times the feature appeared in the top x features across the 100 runs (n). The y-axis is ordered first by n, and ties are broken by the median feature importance, with the top ranking feature appearing at the top of the y-axis.

This feature is highlighted in red, and if the second top ranking feature represented a highly similar cell subset, it was highlighted in pink. We set x = 5 for main figures and trained SCOOP with cancer tissue specific cell features, while in supplementary figures we displayed box plots with x = 10, 5, and 2 and trained SCOOP with cell features across tissues. For all box plots, the center line represents the median, box limits represent the upper and lower quartiles, whiskers represent 1.5x the interquartile range, and dots represent outliers.

### COO prediction

For each cancer type, the most frequent cell subset appearing as the top feature across 100 model runs was predicted as the COO. The COO prediction is consistently highlighted in red (e.g., colon stem cells in Fig. 2b) throughout the manuscript. Except for MPN (Supplementary Fig. 6A), this prediction also corresponded to the most informative feature in the robustness box plots when 5 features remained for all “high confidence” COO predictions (all predictions except those in Supplementary Fig. 7). For MPN, when using blood and bone marrow features, CD34+ CLPs – as opposed to CD34+ early basophils – appeared most frequently when only 5 features remain in the backward selection (Supplementary Fig. 6A). While they do not represent the exact same cell type, they both support the committed hematopoietic progenitors COO hypothesis (see main text).

The second most frequently appearing cell subset, if representing a highly similar cell subset to the COO, was highlighted in pink.

### Statistical testing

For Figs. 1B and 4C, to assess if the variance explained distribution of the top feature (based on mean variance explained) across 100 runs of the model had a significantly different median compared to the next most important feature, we used a one-sided Mann-Whitney test.

Similarly, to assess if the feature importance distribution of the top robustness box plot feature across 100 runs of the model had a significantly different median compared to the next most important feature, we used a one-sided Mann-Whitney test.

To check if the most frequently appearing top feature showed up as the top feature across 100 runs of the model, significantly more so than the second most frequently appearing top feature, we used a one-sided, exact binomial test. Under the null hypothesis, we would expect each feature to appear ½(x+y) times on average, where x and y are the actual number of times the first and second most frequently appearing top feature actually appeared as the top feature, respectively. In certain cases, the first and second top features corresponded to essentially the same feature (e.g., Fig. 2D, GMP from bone marrow and GMP.Neut from bone marrow). In this case, we compared the combined appearances of the top 2 features to the third top feature. In other cases, where the top 2 features were similar but not coming from the same data set, we compared the appearances of the top feature to the third feature. The exact comparison done is indicated on the bar plots by square brackets.

### Variance explained as a function of sample size

To classify cancers into low, medium, and high TMB, we computed the average number of mutations per WGS sample for each cancer type (Supplementary Table 1). Following this, we segmented the cancer types into these three classifications by employing quantile division. We then picked 2 cancers from each category that we had high confidence COO predictions. Per cancer, for each specific number of WGS samples *n*, we randomly sampled *n* patients 100 times, aggregated the sampled patient data SNV profiles, and ran SCOOP on the sampled aggregated mutation profile, using a different random seed for each run.

### WGS sample SNV profile UMAPs

To visualize WGS mutational density profiles across samples we used Uniform Manifold Approximation and Projection (UMAP) dimensionality reduction as implemented in Seurat^92^ except we considered our feature space to be genomic bins rather than genes. Briefly, we normalized the mutational data using the *NormalizeData* function with the parameter *normalization.method* set to “LogNormalize.” Since we have a total of 2,128 bins only, we used all bins in the projection and did not run *FindVariableFeatures*. We then ran *RunPCA* with *npcs* set to “30.” Finally, we ran *RunUMAP* with *dims* set to “1:15.” In the case of lung, we set *min.dist* to “0.2” instead of “0.3”.

### scATAC-seq data analysis via ArchR

#### Cell type UMAPs

To plot the UMAPs of the scATAC-seq data from various cell types, we used ArchR^93^. Specifically, we ran *addIterativeLSI* with *resolution* “0.2”. Then, for Fig. 2A and Fig. 4A (data from ^32^ and ^31^, respectively), we further ran *addHarmony* due to the presence of batch effects (not present in Fig. 2C). Finally, we ran *addUMAP* with *nNeighbors* set to “30” and *minDist* set to “0.5.”

#### Marker gene score computation and visualization

Expression of cell type marker genes was inferred from chromatin accessibility at a gene’s locus using ArchR’s gene score method^93^. Gene scores were visualized using *plotEmbedding* with *colorBy* set to “GeneScoreMatrix” and *quantCut* set to have a range of “0.01” to “0.95” prior to imputation using *addImputeWeights* (Supplementary Fig. 6B).^93^

#### Meta-cell correlation analysis

We began by first sampling meta-cells (i.e groups of similar cells) in scATAC-seq data using a K-nearest-neighbor (KNN) approach. Meta-cells were generated by randomly identifying seed cells for which 500 nearest neighbor cells were then selected based on a KNN graph generated in the latent semantic indexing (LSI) space. Meta-cells were allowed to overlap up to 80%. The function used to conduct this analysis is a customized version of the ArchR function *addCoAccessibility*.

After obtaining meta-cells, we summed the scATAC-seq fragment counts across cells for that meta-cell, representing an aggregated fragment profile. We then correlated all meta-cell profiles with the cancer mutation profile of interest using Pearson’s correlation coefficient. Finally, for each individual cell (i.e. not meta-cell), we assigned it a correlation score corresponding to the mean correlation of the meta-cells it was assigned to. We then plotted the correlation on the scATAC-seq data UMAP per cell. We note that for Fig. 2A, we filtered out cells with a fragment count lower than 10,000 before running any of the previous steps, since we noticed that the meta-cell correlation for that particular dataset correlated with the number of fragments when the fragment count was too low.

### Proliferation rate quantification using scRNA-seq data

To identify the percentage of lung epithelial cells that are proliferating/cycling at homeostasis, we used scRNA-seq data from healthy lung donors^33^. Seurat’s^92^ *CellCycleScoring* function was used to compute the module score for the expression of genes linked to either G1/S or G2/M phase of the cell cycle^94^. Cells with either a G1/S- or G2/M-score greater than 0.1 were classified as cycling and all other cells were considered non-cycling. The percentage of cycling cells for each lung epithelial cell was displayed in Fig. 1F, where cell types were ordered from most to least proliferative.

## ACKNOWLEDGMENTS

This project was funded by the Chan Zuckerberg Initiative (CZI) Data Insights Grant 2022-249299. We thank all members of the Tsankov lab for their feedback.

## AUTHOR CONTRIBUTIONS

P.P. and A.M.T. conceptualized the project. M.D.B., W.L., P.S., B.G., and R.K. assembled and processed the data. M.D.B., B.G., W.L., and A.M.T. performed formal analyses. M.D.B., B.G., W.L. and A.M.T. wrote the manuscript with feedback from P.P., R.K., D.H., E.W., P.S. and approval from all authors.

## Notes

### Competing Interest Statement

The authors have declared no competing interest.

### Summary of Updates

Make sure the abstract on the webpage and PDF correspond to each other. Update the introduction and discussion.

