## Supplementary Figures for "Learning the cellular origins of cancer using single-cell chromatin landscapes"

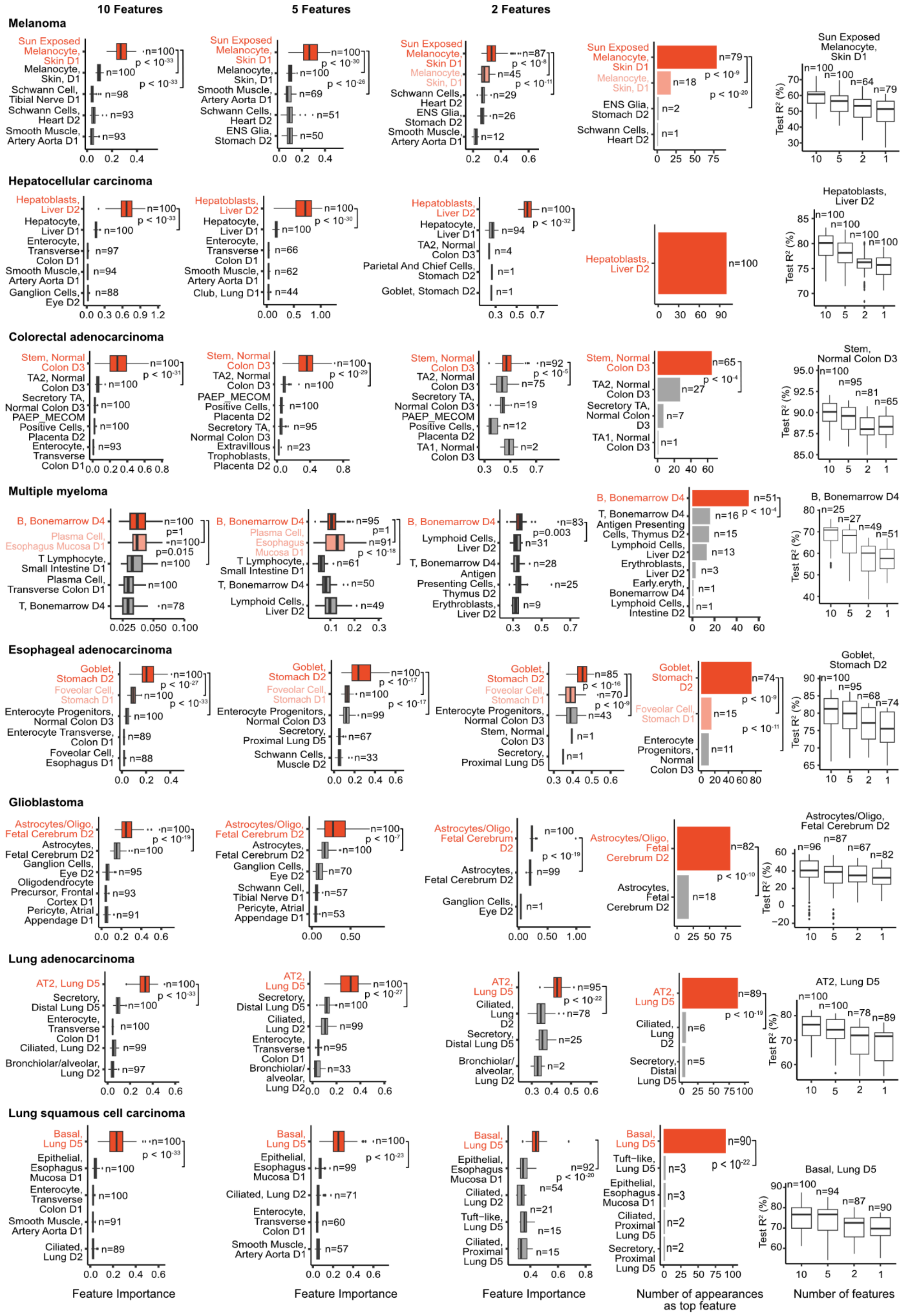

10 Features
5 Features
2 Features
Melanoma
Sun Exposed Melanocyte, Skin D1
Melanocyte, Skin, D1
Schwann Cell, Tibial Nerve D1
Schwann Cells, Heart D2
Smooth Muscle, Artery Aorta D1
Smooth Muscle, Artery Aorta D1
ENS Glia, Stomach D2
ENS Glia, Stomach D2
Sun Exposed Melanocyte, Skin D1
Test R² (%)
Hepatocellular carcinoma
Hepatoblasts, Liver D2
Hepatocyte, Liver D1
Enterocyte, Transverse Colon D1
Smooth Muscle, Artery Aorta D1
Ganglion Cells, Eye D2
Club, Lung D1
TA2, Normal Colon D3
Parietal And Chief Cells, Stomach D2
Goblet, Stomach D2
Colorectal adenocarcinoma
Stem, Normal Colon D3
TA2, Normal Colon D3
Secretory TA, Normal Colon D3
PAEP_MECOM Positive Cells, Placenta D2
Enterocyte, Transverse Colon D1
Extravillous Trophoblasts, Placenta D2
TA1, Normal Colon D3
Multiple myeloma
B, Bonemarrow D4
Plasma Cell, Esophagus Mucosa D1
T Lymphocyte, Small Intestine D1
Plasma Cell, Transverse Colon D1
T, Bonemarrow D4
Lymphoid Cells, Liver D2
Antigen Presenting Cells, Thymus D2
Erythroblasts, Liver D2
Early.eryth, Bonemarrow D4
Lymphoid Cells, Intestine D2
Esophageal adenocarcinoma
Goblet, Stomach D2
Foveolar Cell, Stomach D1
Enterocyte Progenitors, Normal Colon D3
Enterocyte Transverse, Colon D1
Foveolar Cell, Esophagus D1
Secretory, Proximal Lung D5
Schwann Cells, Muscle D2
Stem, Normal Colon D3
Glioblastoma
Astrocytes/Oligo, Fetal Cerebrum D2
Astrocytes, Fetal Cerebrum D2
Ganglion Cells, Eye D2
Oligodendrocyte Precursor, Frontal Cortex D1
Pericyte, Atrial Appendage D1
Schwann Cell, Tibial Nerve D1
Lung adenocarcinoma
AT2, Lung D5
Secretory, Distal Lung D5
Enterocyte, Transverse Colon D1
Ciliated, Lung D2
Bronchiolar/alveolar, Lung D2
Lung squamous cell carcinoma
Basal, Lung D5
Epithelial, Esophagus Mucosa D1
Enterocyte, Transverse Colon D1
Smooth Muscle, Artery Aorta D1
Ciliated, Lung D2
Tuft-like, Lung D5
Ciliated, Proximal Lung D5
Secretory, Proximal Distal Lung D5
Feature Importance
Feature Importance
Feature Importance
Number of appearances as top feature
Number of features

**Supplementary Fig. 1 supporting Fig. 1: SCOOP predicts COOs at unprecedented scale and resolution. Left:** Box plots showing the feature importance for the cancer types in Fig. 1B across 100 SCOOP runs trained on all scATAC-seq cell features with at least 100 cells from [9,10,12–15], where each cell subset is followed by a dataset indicator for that cell subset: D1 for [9], D2 for [10], D3 for [12], D4 for [14], D5 for [13], D6 for [15]. Box plots display the top 5 features when 10, 5, and 2 features remain following backward feature selection (Methods). Highlighted in red are the COOs predicted by SCOOP, in pink are similar cell subsets to the predicted COOs. Also displayed is the number of times the feature appeared in the top 10, 5, and 2 features across the 100 runs (n). The y-axis is ordered first by n, and ties are broken by the median feature importance, with the top ranking feature appearing at the top of the y-axis. Mann-Whitney test *p*-values are displayed, with Bonferroni correction for multiple hypothesis testing when more than one comparison was made. **Middle:** Barplots of the number of times cell subsets appeared as the top feature across 100 SCOOP runs with the most frequently appearing feature (i.e SCOOP's COO prediction) highlighted in red. Exact binomial test p-values are shown (Methods), with Bonferroni correction for multiple hypothesis testing when more than one comparison was made. **Right:** Box plots displaying the test set variance explained (test $R^2$) by the model when 10, 5, 2, and 1 features remained following backward feature selection, and the number of times (n) the top feature at these various iterations of backward feature selection corresponded to the eventual COO prediction. Cell type abbreviations are listed in Supplementary Table 2. For all box plots, the center line represents the median, box limits represent the upper and lower quartiles, whiskers represent 1.5x the interquartile range, and dots represent outliers.

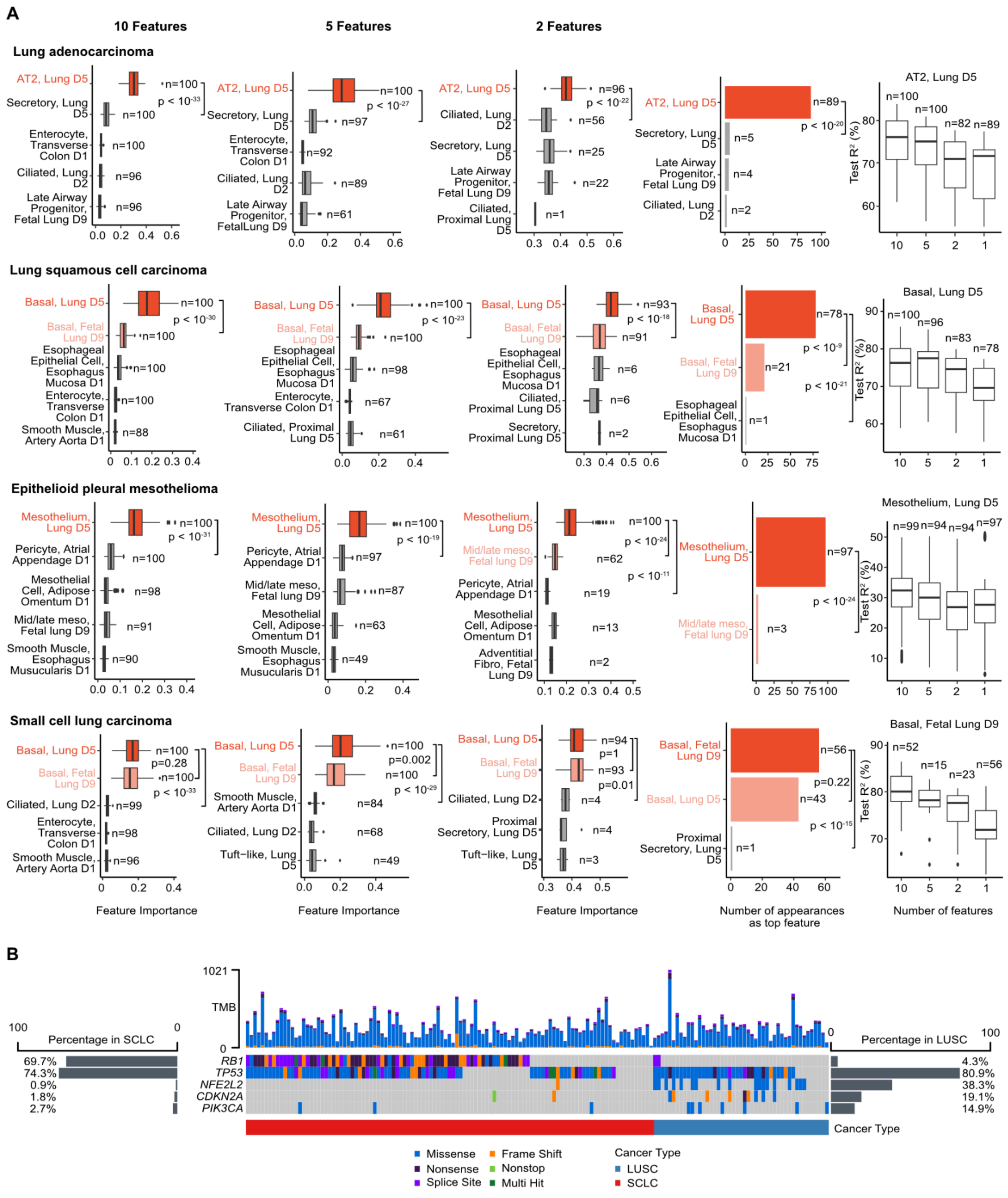


**Supplementary Fig. 2 supporting Fig. 1: Lung cancers are associated with distinct COOs.**

**A) Left:** Box plots showing the feature importance for the cancer types in Fig. 1C across 100 SCOOP runs trained on all scATAC-seq cell features with at least 100 cells from [9,10,12–15,28], where each cell subset is followed by a dataset indicator for that cell subset: D1 for [9], D2 for [10], D3 for [12], D4 for [14], D5 for [13], D6 for [15], D9 for [28]. All cell subsets from the latter datasets were used, with a cell type filter of 100, except for lung neuroendocrine cells from [13], which were also included. Box plots display the top 5 features when 10, 5, and 2 features remain following backward feature selection (Methods). Highlighted in red are the COOs predicted by SCOOP, in pink are similar cell subsets to the predicted COOs. Mann-Whitney test p-values are displayed, with Bonferroni correction for multiple hypothesis testing when more than one comparison was made. **Middle:** Barplots of the number of times cell subsets appeared as the top feature across 100 runs of SCOOP with the most frequently appearing feature (i.e SCOOP's prediction) highlighted in red. Exact binomial test p-values are shown (Methods), with Bonferroni correction for multiple hypothesis testing when more than one comparison was made. **Right:** Box plots displaying the test set variance explained (test $R^2$) by the model when 10, 5, 2, and 1 features remained following backward feature selection, and the top feature at these various iterations of backward feature selection corresponded to the cell type that would eventually be predicted as the COO. **B)** Oncoplot displaying the nonsynonymous somatic mutations for *RB1, TP53, NFE2L2, CDKN2A,* and *PIK3CA* in SCLC and LUSC WGS samples used in our study. The bar plots on the sides show the percentage of mutated samples for each gene in SCLC (left) and LUSC (right). The top bar plot displays the tumor mutation burden (TMB) for each individual sample. Cell type abbreviations are listed in Supplementary Table 2. For all box plots, the center line represents the median, box limits represent the upper and lower quartiles, whiskers represent 1.5x the interquartile range, and dots represent outliers.

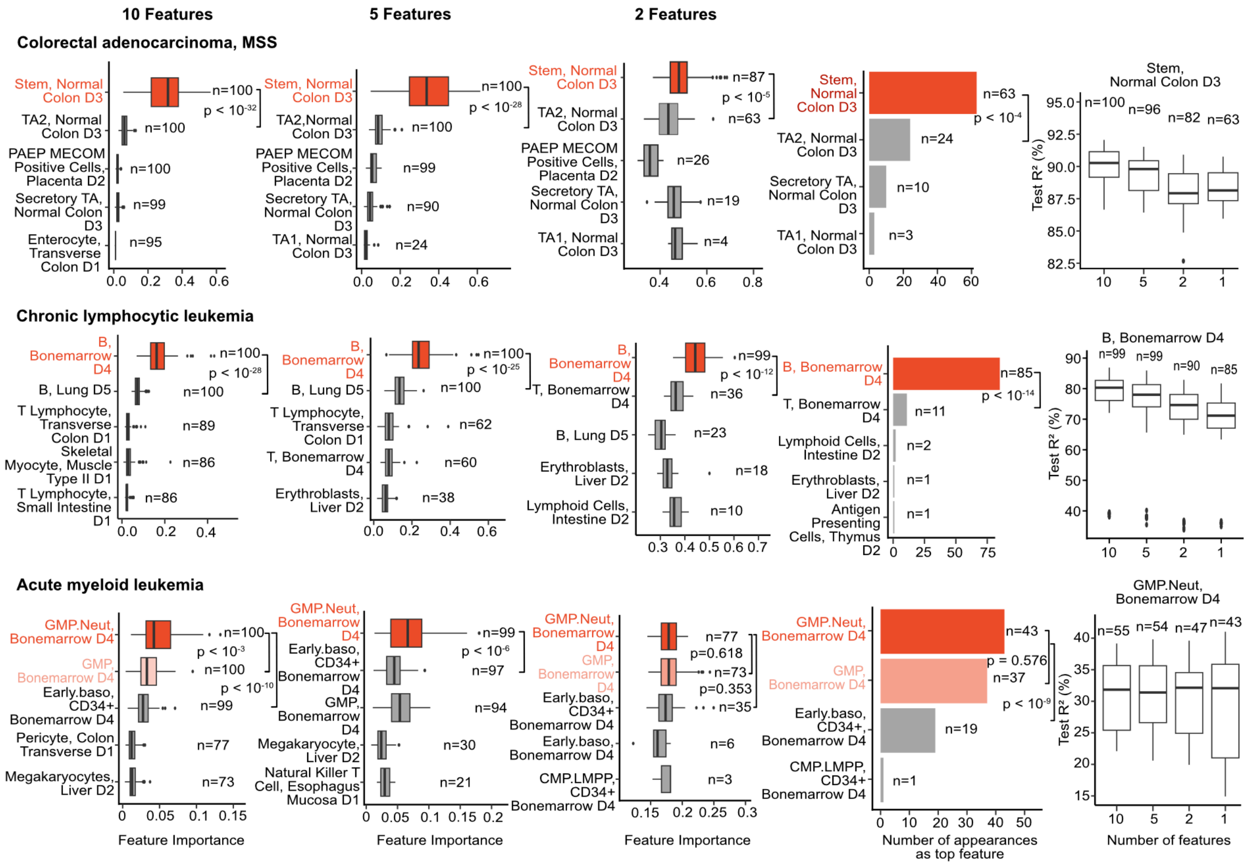


**Supplementary Fig. 3 supporting Fig. 2: SCOOP can pinpoint cell subsets that likely give rise to different tumors. Left:** Box plots showing the feature importance for the cancer types in Fig. 2 across 100 SCOOP runs trained on all scATAC-seq cell features with at least 100 cells from [9,10,12–15], where each cell subset is followed by a dataset indicator for that cell subset: D1 for [9], D2 for [10], D3 for [12], D4 for [14], D5 for [13], D6 for [15]. Box plots display the top 5 features when 10, 5, and 2 features remain following backward feature selection (Methods). Highlighted in red are the COOs predicted by SCOOP, in pink are similar cell subsets to the predicted COOs. Also displayed is the number of times the feature appeared in the top 10, 5, and 2 features across the 100 runs (n). The y-axis is ordered first by n, and ties are broken by the median feature importance, with the top ranking feature appearing at the top of the y-axis. Mann-Whitney test *p*-values are displayed, with Bonferroni correction for multiple hypothesis testing when more than one comparison was made. **Middle:** Barplots of the number of times cell subsets appeared as the top feature across 100 runs of SCOOP with the most frequently appearing feature (i.e SCOOP's prediction) highlighted in red. Exact binomial test p-values are shown (Methods), with Bonferroni correction for multiple hypothesis testing when more than one comparison was made. **Right:** Box plots displaying the test set variance explained (test $R^2$) by the model when 10, 5, 2, and 1 features remained following backward feature selection, and the top feature at these various iterations of backward feature selection corresponded to the cell type that would eventually be predicted as the COO. Cell type abbreviations are listed in Supplementary Table 2. For all box plots, the center line represents the median, box limits represent the upper and lower quartiles, whiskers represent 1.5x the interquartile range, and dots represent outliers.

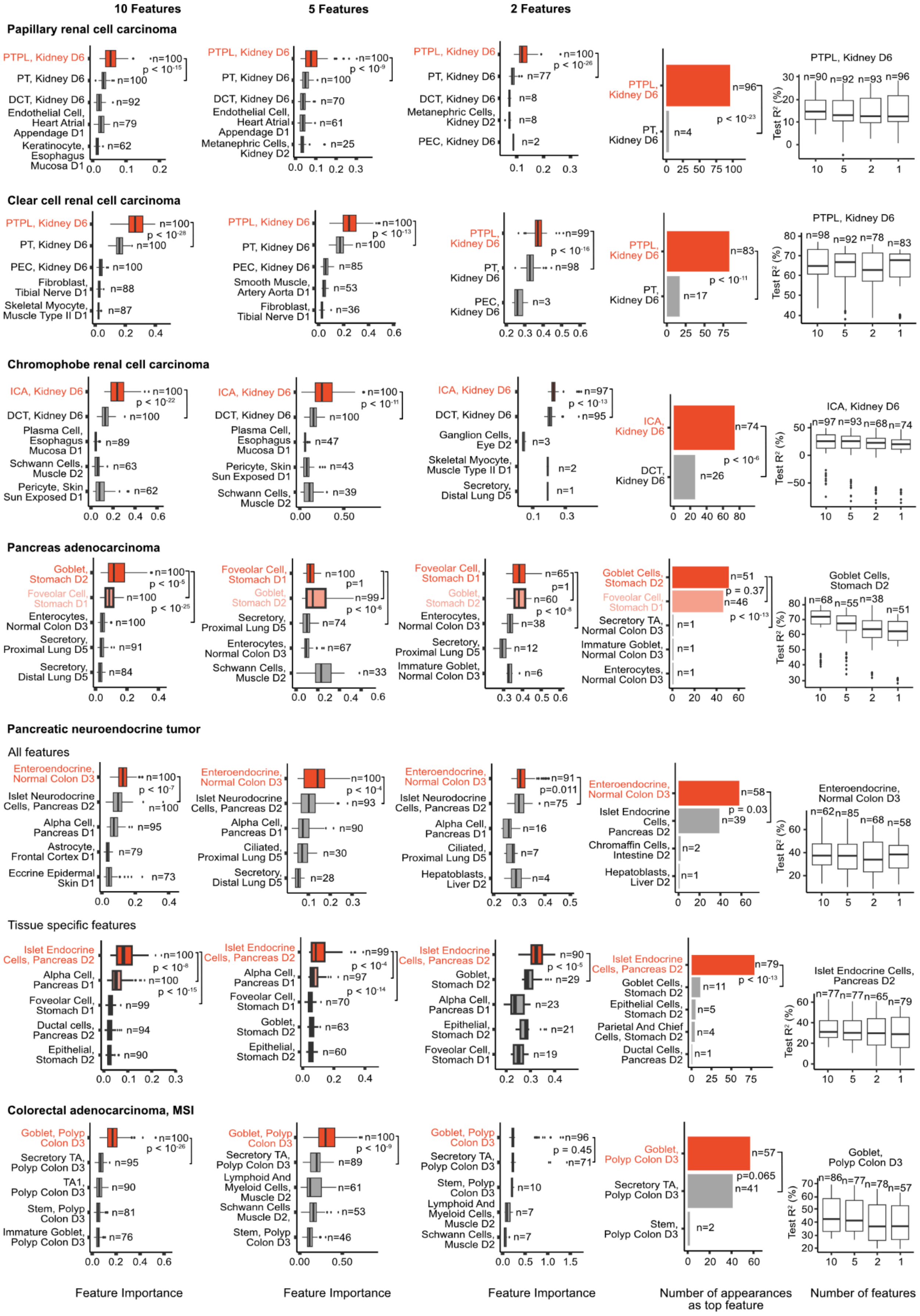
10 Features
5 Features
2 Features
Papillary renal cell carcinoma
PTPL, Kidney D6
PT, Kidney D6
DCT, Kidney D6
Endothelial Cell, Heart Atrial Appendage D1
Keratinocyte, Esophagus Mucosa D1
n=100
p < 10-15
n=92
n=79
n=62
Metanephric Cells, Kidney D2
n=70
n=61
n=25
p < 10-9
PEC, Kidney D6
n=77
n=8
n=2
p < 10-26
PTPL, Kidney D6
PT, Kidney D6
n=96
n=4
p < 10-23
Test R² (%)
n=90 n=92 n=93 n=96
Clear cell renal cell carcinoma
PEC, Kidney D6
Fibroblast, Tibial Nerve D1
Skeletal Myocyte, Muscle Type II D1
p < 10-28
n=88
n=87
Smooth Muscle, Artery Aorta D1
n=85
n=53
n=36
p < 10-13
n=99
n=98
n=3
p < 10-16
n=83
n=17
p < 10-11
n=98 n=92 n=78 n=83
Chromophobe renal cell carcinoma
ICA, Kidney D6
Plasma Cell, Esophagus Mucosa D1
Schwann Cells, Muscle D2
Pericyte, Skin Sun Exposed D1
p < 10-22
n=89
n=63
n=62
n=47
n=43
n=39
p < 10-11
Ganglion Cells, Eye D2
Skeletal Myocyte, Muscle Type II D1
Secretory, Distal Lung D5
n=97
n=95
p < 10-13
n=1
ICA, Kidney D6
DCT, Kidney D6
n=74
n=26
p < 10-6
n=97 n=93 n=68 n=74
Pancreas adenocarcinoma
Goblet, Stomach D2
Foveolar Cell, Stomach D1
Enterocytes, Normal Colon D3
Secretory, Proximal Lung D5
Secretory, Distal Lung D5
p < 10-5
p < 10-25
n=91
n=84
p=1
p < 10-6
n=74
n=67
n=33
n=65
n=60
p < 10-8
n=38
n=12
Immature Goblet, Normal Colon D3
n=6
Goblet Cells, Stomach D2
Foveolar Cell, Stomach D1
Secretory TA, Normal Colon D3
n=51
n=46
p = 0.37
p < 10-13
Goblet Cells, Stomach D2
n=68 n=55 n=38 n=51
Pancreatic neuroendocrine tumor
All features
Enteroendocrine, Normal Colon D3
Islet Neurodocrine Cells, Pancreas D2
Alpha Cell, Pancreas D1
Astrocyte, Frontal Cortex D1
Eccrine Epidermal Skin D1
p < 10-7
n=95
n=73
Ciliated, Proximal Lung D5
n=93
n=90
n=30
n=28
p < 10-4
Hepatoblasts, Liver D2
n=91
p=0.011
n=75
n=16
n=7
Enteroendocrine, Normal Colon D3
Islet Endocrine Cells, Pancreas D2
Chromaffin Cells, Intestine D2
n=58
n=39
p = 0.03
n=62 n=85 n=68 n=58
Tissue specific features
Islet Endocrine Cells, Pancreas D2
Ductal cells, Pancreas D2
Epithelial, Stomach D2
p < 10-8
p < 10-15
n=94
n=97
p < 10-14
n=70
n=63
n=60
Goblet, Stomach D2
p < 10-5
n=29
n=23
n=21
n=19
Goblet Cells, Stomach D2
Epithelial Cells, Stomach D2
Parietal And Chief Cells, Stomach D2
n=79
n=11
n=5
p < 10-13
Islet Endocrine Cells, Pancreas D2
n=77 n=77 n=65 n=79
Colorectal adenocarcinoma, MSI
Goblet, Polyp Colon D3
Secretory TA, Polyp Colon D3
TA1, Polyp Colon D3
Stem, Polyp Colon D3
Immature Goblet, Polyp Colon D3
p < 10-26
n=81
n=76
Lymphoid And Myeloid Cells, Muscle D2
Schwann Cells, Muscle D2,
n=89
n=61
n=46
p < 10-9
p = 0.45
n=71
n=10
Goblet, Polyp Colon D3
n=57
n=41
p=0.065
n=86 n=77 n=78 n=57
Feature Importance
Number of appearances as top feature
Number of features

**Supplementary Fig. 4 supporting Fig. 3: Histological cancer subtypes are associated with different COOs. Left:** Box plots showing the feature importance for the cancer types in Fig. 3 across 100 SCOOP runs trained on all scATAC-seq cell features with at least 100 cells from [9,10,12–15], where each cell subset is followed by a dataset indicator for that cell subset: D1 for [9], D2 for [10], D3 for [12], D4 for [14], D5 for [13], D6 for [15]. Box plots display the top 5 features when 10, 5, and 2 features remain following backward feature selection (Methods). Highlighted in red are the COOs predicted by SCOOP, in pink are similar cell subsets to the predicted COOs. Also displayed is the number of times the feature appeared in the top 10, 5, and 2 features across the 100 runs (n). The y-axis is ordered first by n, and ties are broken by the median feature importance, with the top ranking feature appearing at the top of the y-axis. Mann-Whitney test *p*-values are displayed, with Bonferroni correction for multiple hypothesis testing when more than one comparison was made. **Middle:** Barplots of the number of times cell subsets appeared as the top feature across 100 runs of SCOOP with the most frequently appearing feature (i.e SCOOP's prediction) highlighted in red. Exact binomial test p-values are shown (Methods), with Bonferroni correction for multiple hypothesis testing when more than one comparison was made. **Right:** Box plots displaying the test set variance explained (test $R^2$) by the model when 10, 5, 2, and 1 features remained following backward feature selection, and the top feature at these various iterations of backward feature selection corresponded to the cell type that would eventually be predicted as the COO. Cell type abbreviations are listed in Supplementary Table 2. For all box plots, the center line represents the median, box limits represent the upper and lower quartiles, whiskers represent 1.5x the interquartile range, and dots represent outliers.

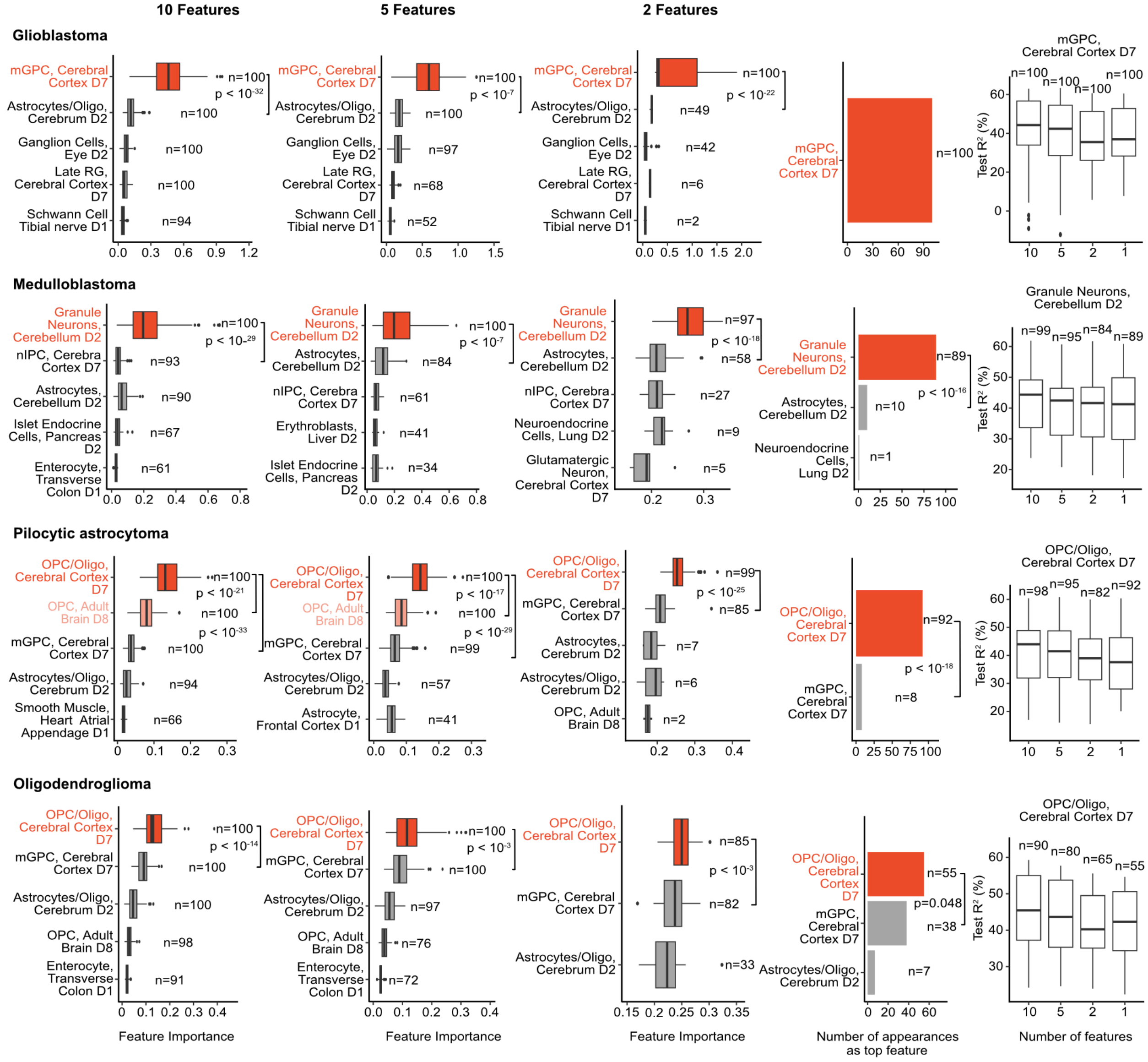


**Supplementary Fig. 5 supporting Fig. 4: Gliomas likely arise from fetal-like multipotent progenitor cells. Left:** Box plots showing the feature importance for the cancer types in Fig. 4 across 100 SCOOP runs trained on all scATAC-seq cell features with at least 100 cells from [9–15,46], where each cell subset is followed by a dataset indicator for that cell subset: D1 for [9], D2 for [10], D3 for [12], D4 for [14], D5 for [13], D6 for [15]D7 for [11] , D8 for [46]. Box plots display the top 5 features when 10, 5, and 2 features remain following backward feature selection (Methods). Highlighted in red are the COOs predicted by SCOOP, in pink are similar cell subsets to the predicted COOs. Also displayed is the number of times the feature appeared in the top 10, 5, and

2 features across the 100 runs (n). The y-axis is ordered first by n, and ties are broken by the median feature importance, with the top ranking feature appearing at the top of the y-axis. Mann-Whitney test *p*-values are displayed, with Bonferroni correction for multiple hypothesis testing when more than one comparison was made. **Middle:** Barplots of the number of times cell subsets appeared as the top feature across 100 runs of SCOOP with the most frequently appearing feature (i.e SCOOP's prediction) highlighted in red. Exact binomial test p-values are shown (Methods), with Bonferroni correction for multiple hypothesis testing when more than one comparison was made. **Right:** Box plots displaying the test set variance explained (test $R^2$) by the model when 10, 5, 2, and 1 features remained following backward feature selection, and the top feature at these various iterations of backward feature selection corresponded to the cell type that would eventually be predicted as the COO. Cell type abbreviations are listed in Supplementary Table 2. For all box plots, the center line represents the median, box limits represent the upper and lower quartiles, whiskers represent 1.5x the interquartile range, and dots represent outliers.

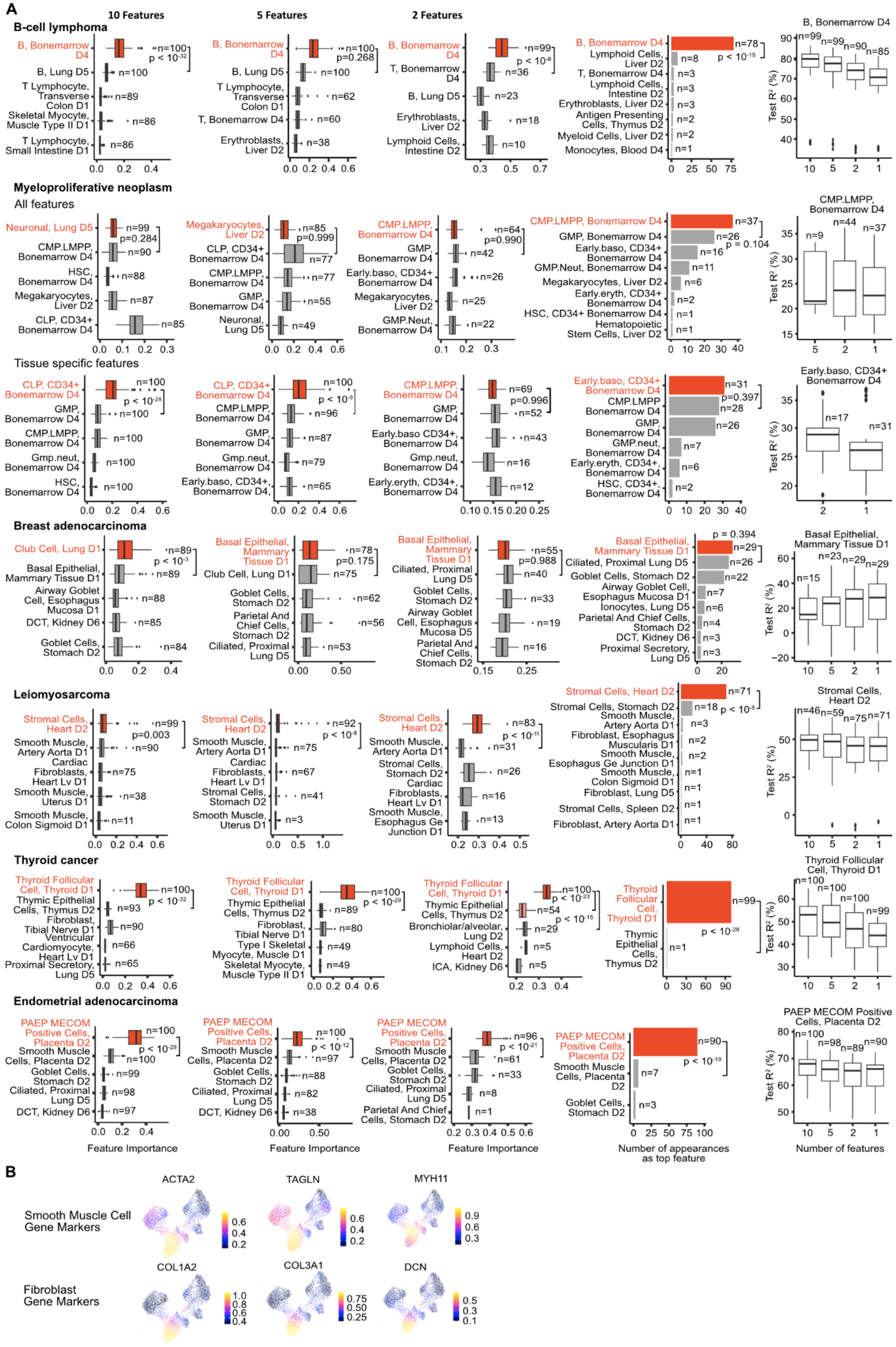

A
10 Features
5 Features
2 Features
B-cell lymphoma
B, Bonemarrow D4
B, Lung D5
T Lymphocyte, Transverse Colon D1
Skeletal Myocyte, Muscle Type II D1
T Lymphocyte, Small Intestine D1
T, Bonemarrow D4
Erythroblasts, Liver D2
Lymphoid Cells, Intestine D2
Antigen Presenting Cells, Thymus D2
Myeloid Cells, Liver D2
Monocytes, Blood D4
Myeloproliferative neoplasm
All features
Neuronal, Lung D5
CMP.LMPP, Bonemarrow D4
HSC, Bonemarrow D4
Megakaryocytes, Liver D2
CLP, CD34+ Bonemarrow D4
GMP, Bonemarrow D4
Early.baso, CD34+ Bonemarrow D4
GMP.Neut, Bonemarrow D4
Early.eryth, CD34+ Bonemarrow D4
Hematopoietic Stem Cells, Liver D2
Tissue specific features
Gmp.neut, Bonemarrow D4
Breast adenocarcinoma
Club Cell, Lung D1
Basal Epithelial, Mammary Tissue D1
Airway Goblet Cell, Esophagus Mucosa D1
DCT, Kidney D6
Goblet Cells, Stomach D2
Parietal And Chief Cells, Stomach D2
Ciliated, Proximal Lung D5
Ionocytes, Lung D5
Proximal Secretory, Lung D5
Leiomyosarcoma
Stromal Cells, Heart D2
Smooth Muscle, Artery Aorta D1
Cardiac Fibroblasts, Heart Lv D1
Smooth Muscle, Uterus D1
Smooth Muscle, Colon Sigmoid D1
Stromal Cells, Stomach D2
Smooth Muscle, Esophagus Ge Junction D1
Fibroblast, Esophagus Muscularis D1
Fibroblast, Lung D5
Stromal Cells, Spleen D2
Fibroblast, Artery Aorta D1
Thyroid cancer
Thyroid Follicular Cell, Thyroid D1
Thymic Epithelial Cells, Thymus D2
Fibroblast, Tibial Nerve D1
Ventricular Cardiomyocyte, Heart Lv D1
Proximal Secretory, Lung D5
Type I Skeletal Myocyte, Muscle D1
Skeletal Myocyte, Muscle Type II D1
Bronchiolar/alveolar, Lung D2
Lymphoid Cells, Heart D2
ICA, Kidney D6
Endometrial adenocarcinoma
PAEP MECOM Positive Cells, Placenta D2
Smooth Muscle Cells, Placenta D2
Goblet Cells, Stomach D2
Ciliated, Proximal Lung D5
DCT, Kidney D6
Parietal And Chief Cells, Stomach D2
Feature Importance
Number of appearances as top feature
Number of features
Test R² (%)
B
Smooth Muscle Cell Gene Markers
ACTA2
TAGLN
MYH11
Fibroblast Gene Markers
COL1A2
COL3A1
DCN

**Supplementary Fig. 6 supporting Fig. 4: Pan-cancer COO predictions identifies metaplasia-mediated neoplasms. A) Left:** Box plots showing the feature importance for the cancer types in Fig. 3 across 100 SCOOP runs trained on all scATAC-seq cell features with at least 100 cells from [9,10,12–15], where each cell subset is followed by a dataset indicator for that cell subset: D1 for [9], D2 for [10], D3 for [12], D4 for [14], D5 for [13], D6 for [15]. Box plots display the top 5 features when 10, 5, and 2 features remain following backward feature selection (Methods). Highlighted in red are the COOs predicted by SCOOP, in pink are similar cell subsets to the predicted COOs. Also displayed is the number of times the feature appeared in the top 10, 5, and 2 features across the 100 runs (n). The y-axis is ordered first by n, and ties are broken by the median feature importance, with the top ranking feature appearing at the top of the y-axis. Mann-Whitney test *p*-values are displayed, with Bonferroni correction for multiple hypothesis testing when more than one comparison was made. **Middle:** Barplots of the number of times cell subsets appeared as the top feature across 100 runs of SCOOP with the most frequently appearing feature (i.e SCOOP's prediction) highlighted in red. Exact binomial test p-values are shown (Methods), with Bonferroni correction for multiple hypothesis testing when more than one comparison was made. **Right:** Box plots displaying the test set variance explained (test $R^2$) by the model when 10, 5, 2, and 1 features remained following backward feature selection, and the top feature at these various iterations of backward feature selection corresponded to the cell type that would eventually be predicted as the COO. **B)** UMAP supporting Leiomyosarcoma COO prediction of stromal cells, showing that this cell population exhibits high chromatin accessibility at the locus of various smooth muscle cell (ACTA2, TAGLN, MYH11; expected COO) and fibroblast (COL1A2, COL3A1, DCN) markers as calculated by ArchR (Methods). Cell type abbreviations are listed in Supplementary Table 2. For all box plots, the center line represents the median, box limits represent the upper and lower quartiles, whiskers represent 1.5x the interquartile range, and dots represent outliers.

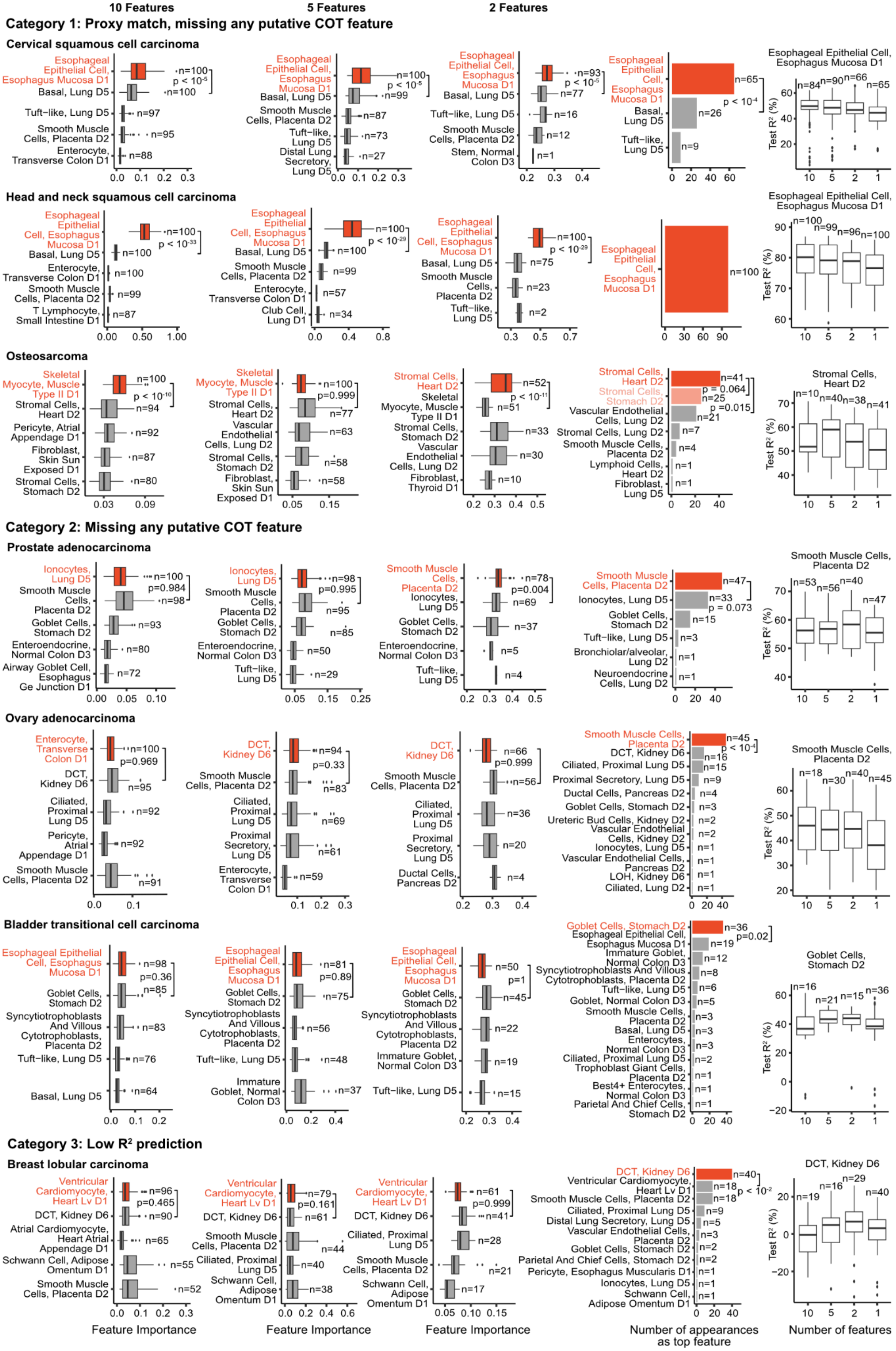
10 Features
5 Features
2 Features
Category 1: Proxy match, missing any putative COT feature
Cervical squamous cell carcinoma
Esophageal Epithelial Cell, Esophagus Mucosa D1
Basal, Lung D5
Tuft−like, Lung D5
Smooth Muscle Cells, Placenta D2
Enterocyte, Transverse Colon D1
Distal Lung Secretory, Lung D5
Stem, Normal Colon D3
Esophageal Epithelial Cell, Esophagus Mucosa D1
Head and neck squamous cell carcinoma
Enterocyte, Transverse Colon D1
T Lymphocyte, Small Intestine D1
Club Cell, Lung D1
Osteosarcoma
Skeletal Myocyte, Muscle Type II D1
Stromal Cells, Heart D2
Pericyte, Atrial Appendage D1
Fibroblast, Skin Sun Exposed D1
Stromal Cells, Stomach D2
Vascular Endothelial Cells, Lung D2
Fibroblast, Thyroid D1
Smooth Muscle Cells, Placenta D2
Lymphoid Cells, Heart D2
Fibroblast, Lung D5
Category 2: Missing any putative COT feature
Prostate adenocarcinoma
Ionocytes, Lung D5
Goblet Cells, Stomach D2
Enteroendocrine, Normal Colon D3
Airway Goblet Cell, Esophagus Ge Junction D1
Bronchiolar/alveolar, Lung D2
Neuroendocrine Cells, Lung D2
Ovary adenocarcinoma
Enterocyte, Transverse Colon D1
DCT, Kidney D6
Ciliated, Proximal Lung D5
Pericyte, Atrial Appendage D1
Proximal Secretory, Lung D5
Ductal Cells, Pancreas D2
Ureteric Bud Cells, Kidney D2
Vascular Endothelial Cells, Kidney D2
Vascular Endothelial Cells, Pancreas D2
LOH, Kidney D6
Ciliated, Lung D2
Bladder transitional cell carcinoma
Syncytiotrophoblasts And Villous Cytotrophoblasts, Placenta D2
Immature Goblet, Normal Colon D3
Goblet, Normal Colon D3
Trophoblast Giant Cells, Placenta D2
Best4+ Enterocytes, Normal Colon D3
Parietal And Chief Cells, Stomach D2
Category 3: Low $R^2$ prediction
Breast lobular carcinoma
Ventricular Cardiomyocyte, Heart Lv D1
Atrial Cardiomyocyte, Heart Atrial Appendage D1
Schwann Cell, Adipose Omentum D1
Pericyte, Esophagus Muscularis D1
Feature Importance
Number of appearances as top feature
Test $R^2$ (%)
Number of features

**Supplementary Fig. 7: SCOOP COO predictions on additional cancers highlight method limitations. Left:** Box plots showing the feature importance for category 1, 2, and 3 COO predictions across 100 SCOOP runs trained on all scATAC-seq cell features with at least 100 cells from [9,10,12–15], where each cell subset is followed by a dataset indicator for that cell subset: D1 for [9], D2 for [10], D3 for [12], D4 for [14], D5 for [13], D6 for [15]. Box plots display the top 5 features when 10, 5, and 2 features remain following backward feature selection (Methods). Highlighted in red are the COOs predicted by SCOOP, in pink are similar cell subsets to the predicted COOs. Also displayed is the number of times the feature appeared in the top 10, 5, and 2 features across the 100 runs (n). The y-axis is ordered first by n, and ties are broken by the median feature importance, with the top ranking feature appearing at the top of the y-axis. Mann-Whitney test *p*-values are displayed, with Bonferroni correction for multiple hypothesis testing when more than one comparison was made. **Middle:** Barplots of the number of times cell subsets appeared as the top feature across 100 runs of SCOOP with the most frequently appearing feature (i.e SCOOP's prediction) highlighted in red. Exact binomial test p-values are shown (Methods), with Bonferroni correction for multiple hypothesis testing when more than one comparison was made. **Right**: Box plots displaying the test set variance explained (test $R^2$) by the model when 10, 5, 2, and 1 features remained following backward feature selection, and the top feature at these various iterations of backward feature selection corresponded to the cell type that would eventually be predicted as the COO. Cell type abbreviations are listed in Supplementary Table 2. For all box plots, the center line represents the median, box limits represent the upper and lower quartiles, whiskers represent 1.5x the interquartile range, and dots represent outliers.
